# Development and validation of the Irish Science Self-Efficacy Children’s Questionnaire to assess the short-term influence of scientists facilitating outreach

**DOI:** 10.1101/2020.01.25.919357

**Authors:** Sarah Carroll, Jerome Sheahan, Veronica McCauley, Muriel Grenon

**Author notes:** Corresponding Authors E-mail addresses (MG) and (SC).

## Abstract

Despite the positive relationship between science self-efficacy and science motivation, and the prevalence of scientist-led outreach programs, there is limited research on the effect of scientists on children’s science self-efficacy beliefs. Furthermore, there is no extant science self-efficacy research on primary pupils in Ireland, possibly due to the absence of a suitable and valid assessment instrument. This multi-study research aimed to develop and validate a questionnaire suitable for investigating the effect of scientist-facilitated science outreach on the science self-efficacy of children aged 11-12 years old. In Study 1 the first version of the Irish Science Self-Efficacy Children’s Questionnaire (IS-SEC-Q) was developed and tested with 92 primary school students. In Study 2, the revised questionnaire was re-tested with 282 students. For both studies, construct validity was examined through factor analysis. Questionnaire interpretation and comprehension were investigated via interviews (N=4 and 25 respectively). In Study 2, convergent and criterion validity of the scales was also examined. The final IS-SEC-Q contained 5 scales (63 items), including an adaptation of Usher and Pajares ‘Sources of Self-Efficacy in Mathematics’ scale. Study 2 interviews indicated possible misinterpretations of the Emotional State subscale. The questionnaire demonstrated good psychometric properties and should serve well for informal science education practitioners endeavouring to assess their impact on primary children’s science self-efficacy.

## Introduction

To address shortfalls in science graduates and improve scientific literacy, informal science education initiatives aim to increase children’s engagement with science (1–5). Many of these informal science education initiatives include outreach programs where scientists act as facilitators and engage in hands-on science activities with participants (5, 6). This is also becoming more prevalent, as the European Commission has reported a rising trend in scientists participating in both face-to-face interaction and delivering hands-on exploration of science activities with the public (7). In fact, according to a survey of STEM outreach programs in the U.S., the involvement of scientists as ‘role models’ is one of six most common pedagogical features in such science outreach programs (1).

Such informal science education initiatives often focus on improving participant’s engagement with science. Recent research indicates that whilst children are in fact interested and enjoy science (8–10), many children still view science as not “…for them” (8). In addition, young children (10-11 years old) perceive science as being a “hard” subject that requires concentration and motivation (8). Therefore, in their pursuit of improving children’s engagement with science, informal science education initiatives should consider targeting another outcome in addition to increased interest. Such an outcome may be science self-efficacy, which is positively linked to academic performance, motivation and aspiration in science (11–13).

### Science self-efficacy

First postulated by Psychologist Albert Bandura, self-efficacy is an extension of social cognitive theory that can be defined as “… a judgment of one’s ability to organize and execute given types of performances…” (11). In comparison to those with low self-efficacy, individuals with high self-efficacy are more likely to preserver through challenges (12). SSE is task-characterised and centred around objectives individuals can do, as opposed to objectives that individuals will do, which is related to intentions (14).

Science Self-Efficacy (SSE) can be described as an individual’s self confidence in completing tasks successfully in science. These tasks could be performing certain scientific skills or answering science-based questions. Due to its positive correlation with academic performance (13, 15), investigations into how SSE beliefs are formed and influenced have been a popular topic of interest for educational researchers. However, despite the fact that self-efficacy beliefs seem to be at their most malleable in early stages of learning (11), most extant studies have focused on post-primary level education (16–19), with limited research involving primary-level education students (15, 20). In addition, whilst it has been demonstrated that peers, teachers and family members can all positively affect an individual’s academic self-efficacy (21, 22), the influence of scientists, in any capacity, is yet unknown. Due to the increasing prevalence of scientists delivering science public engagement activities to children (7), this should be investigated to recommend best practice for facilitators.

### The four sources of science self-efficacy

Bandura (11) proposed that self-efficacy has 4 determinants, termed ‘sources’, which have since been empirically supported by numerous studies (13, 23). These four sources are: Mastery Experience, Vicarious Experience, Verbal Persuasion and Emotional State (11). Two of these sources, Mastery Experience and Emotional State, stem from oneself. The other two, Vicarious Experience and Verbal Persuasion, are provided by others. These four sources can be described as follows:

1. Mastery Experience has been repeatedly shown to be the strongest predictor of self-efficacy (11, 13), and can be described as an individual’s past successful performances in completing tasks.
2. Emotional State concerns an individual’s mood when completing a task. A general good mood can raise self-efficacy beliefs and subsequent performance whereas negative feelings, such as stress or sadness, can be perceived as a lack of self-efficacy regarding that task (24).
3. Vicarious Experience regards an individual’s ability towards the completion of a specific task when compared to the attainments of others (11). This is particularly relevant when the individual has not encountered a specific problem before, and therefore has no previous experiences on which to gauge their perceived capabilities. For example, a student who may have received a ‘C’ in an exam, may compare themselves with peers to fully comprehend their result (25). This idea of gauging the experience of others can be referred to as ‘social modelling’. There are two forms of this modelling: ‘coping’ and ‘mastery’. Coping modelling is where individual’s watch their peers initially struggle to complete tasks but then eventually succeed. In this way, the observed performers have coped with the encountered challenge, hence the name. Mastery modelling is when individuals watch others completing a task without any seeming difficulty (11).
4. Verbal Persuasion is when “…significant others express faith in one’s abilities” (11). This is particularly pertinent when expressed at times where the student may be struggling with a task (11). The provision of Verbal Persuasion can be strengthened when accompanied by instructions and conditions that help bring about success. In addition, Verbal Persuasion is most effective when the persuader is deemed credibly competent by the individual performing the task (11, 26). In other words, if the performer believes that the person giving the praise is proficient in that specific task, then they are more willing to accept the praise and process it as being meaningful.

### Science self-efficacy and scientists

Science outreach activities facilitated by scientists may unknowingly be providing all four of these sources. For example, the use of hands-on activities, group-learning (teamwork) and scientists as role models or facilitators are three common pedagogical features of informal science education activities (1, 4). In such a scenario, there may be several ways that participants may be drawing upon sources of science self-efficacy.

Firstly, hands-on activities that involve successfully performing a science experiment or related scientific skills, may provide children opportunities to gain more Mastery Experience. Secondly, if children perceive the scientist facilitator to be credibly competent in the science task at hand, any praise or feedback given by the scientist may serve as Verbal Persuasion. Bandura (11) stated that this combination of Verbal Persuasion and successful examples of Mastery Experience is especially effective. Thirdly, science outreach programs generally tailor their activities to be fun and engaging, which could induce a positive Emotional State (and reduce any existing anxiety relating to that activity). Lastly, there could be several layers of Vicarious Experience at play. Scientist facilitators demonstrating how to perform experimental steps, without any difficulty, may be perceived by children as providing examples of mastery modelling. Such displays of mastery modelling coupled with effective instruction have also been shown to be uniformly effective in increasing self-efficacy (11). Watch their peers performing the same steps may also contribute to received Vicarious Experience, as an example of coping modelling.

To determine what effect scientist facilitators in such science outreach activities may have on children’s SSE beliefs, it is necessary to assess beliefs before and after participation in an outreach session and investigate whether any of the incurred changes are due to an increase in any of the four sources of self-efficacy. There has been a variety of different questionnaires used to assess the strength of science self-efficacy in different contexts (27). However, to be accurate, such instruments must be tailored to their specific research context (27). There have been no published questionnaires assessing the strength of primary pupil’s science self-efficacy as it relates to the Irish primary science curriculum.

For assessing the influence of the four different sources of self-efficacy, many researchers have adapted the ‘Sources of Mathematics Self-Efficacy’s scale; developed by Lent, Lopez, and Bieschke (28), to fit their own research context (16,18,21). For example, Usher & Pajares (29) adapted this scale to create their ‘Sources of Middle School Mathematics Self-Efficacy’ scale. In their subsequent validation paper they suggested that it could be further adapted and validated again for other subjects and educational contexts (30). Their scale comprises items that assess the influence of the four sources of SSE and includes family members and teachers as providers of Verbal Persuasion and Vicarious Experience but does not include scientists.

In science outreach, whilst there have been scales developed to assess the self-efficacy of the scientists delivering the outreach (30–31) currently there are no instruments to assess the impact of the outreach on participating children’s SSE.

In summary, in the existing literature there is an absence of a suitable instrument to assess the influence of scientist-led hands-on outreach on Irish primary pupil’s SSE. To address this, this work describes the development and evaluation of the Irish Science Self-Efficacy Children’s Questionnaire (IS-SEC-Q) as a suitably valid and reliable instrument for this type of investigation. This research had the following objectives:

1. Develop a questionnaire containing scales to assess the strength and sources of 11-12 year old children’s SSE. The questionnaire must:

- relate to the Irish primary science curriculum
- assess the four sources of self-efficacy
- includes scientists as potential providers of Vicarious Experience and Verbal Persuasion
- be suitable to be used before and after participation in a scientist-facilitated hands-on outreach session
2. Evaluate the validity and reliability of the resulting questionnaire by specifically examining questionnaire interpretation, construct-related validity, convergent-related validity, criterion-related validity and internal consistency of the scales.

### Overview of the two studies

The development and validation of the IS-SEC-Q took place over two studies (see Fig 1 for schematic of the developmental strategy). The developmental strategy addresses five key steps of questionnaire development: (a) initial item design/adaptation, (b) expert review of items, (c) pre-testing of items with small sample of intended audience, (d) piloting the items, (e) evaluating the validity and internal consistency of the scales as a whole (32). The purpose of the last step is the ascertain that the questionnaire ‘works’ as intended *i.e.* it accurately and reliably assess the construct(s) of interest. Therefore, the last two steps (piloting and evaluation) can be cyclical, with each testing of the questionnaire informing new changes to improve its psychometric properties. This process continues until the validity of the questionnaire is satisfactory.

**Fig 1.**
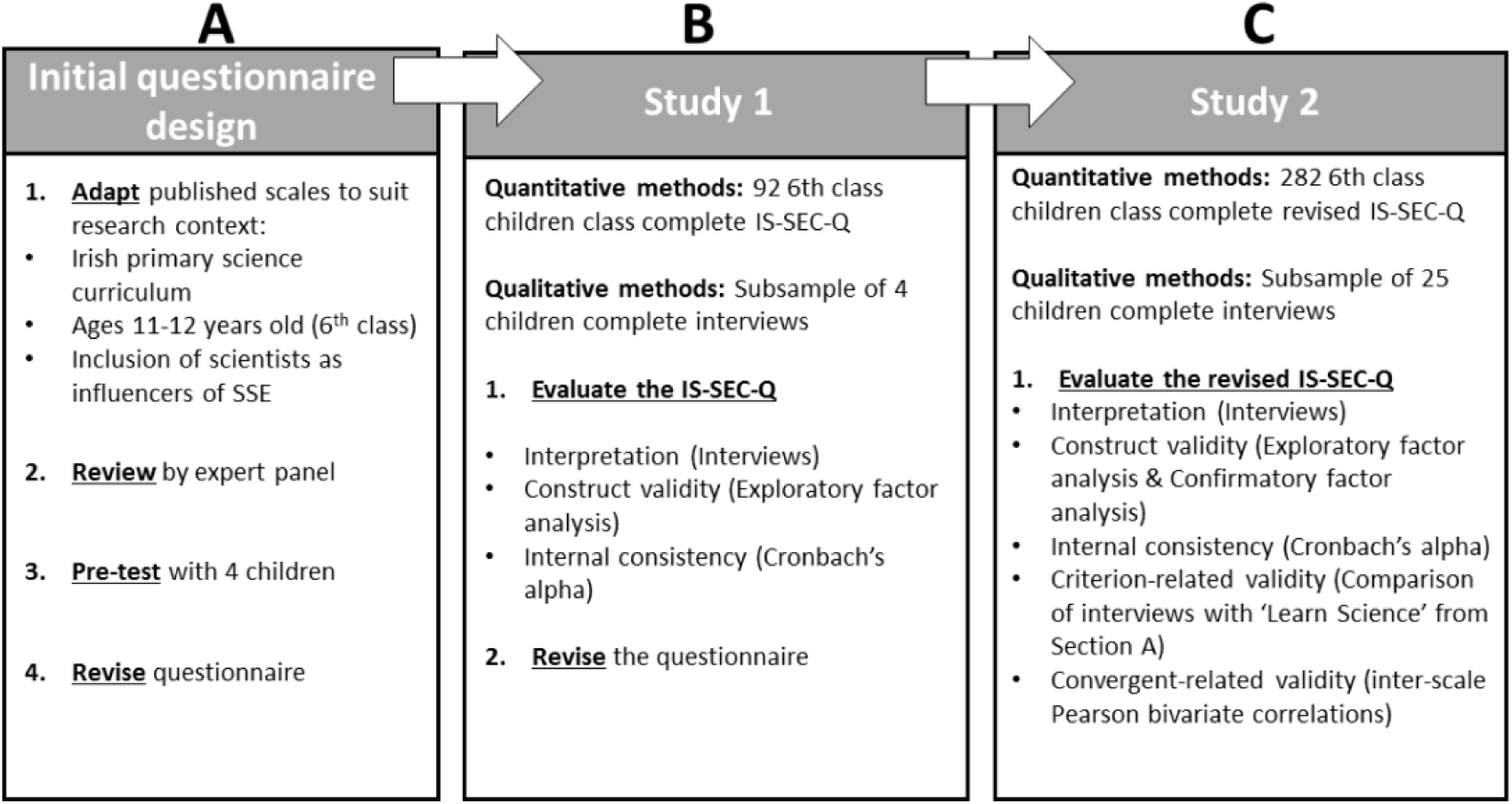
The developmental strategy of the Irish Science Self-Efficacy Children’s Questionnaire (IS-SEC-Q). The development of the questionnaire addressed key steps of scale development as recommended by both Bandura (2006) and DeVellis (2017). The validation of the IS-SEC-Q used two independent datasets in two separate studies

In this work, after the initial questionnaire (S1 Figure) was constructed (A in Fig 1), it was piloted with 92 children (B in Fig 1). This pilot is henceforth referred to as Study 1. In Study 1, the comprehension of the questionnaire and construct validity of the scales was examined. The results of Study 1 indicated that certain changes had to be made to improve the validity of the questionnaire. These changes led to the second version of the IS-SEC-Q (S2 Figure), which was re-tested in Study 2 with a larger sample of 282 children (C in Fig 1). Study 2 examined the construct-, convergent- and criterion-related validity of the questionnaire. In Study 2 there was also a more detailed examination of user comprehension and interpretation.

For both studies, participating primary schools based in county Galway in Ireland were recruited via convenience sampling and included mixed (boys and girls) and single sex (i.e. boys or girls only) schools. The language of instruction of participating schools was either English or Irish (Gaelscoils). Data was only collected from participants from 6th class (11-12 years old) who had guardian consent and child assent. The studies were granted ethical approval by the research ethics committee of the National University of Ireland Galway.

### The initial construction of the Irish Science Self-Efficacy Children’s Questionnaire

The construction of the IS-SEC-Q followed recommendations by Bandura (27) in his ‘Guide for constructing self-efficacy scales’. Following an extensive review of extant self-efficacy instruments, an initial pool of items was generated. The name, purpose, origin, adaptations, and example items of each scale is outlined in Table 1. The resulting questionnaire (S1 Figure) contained five scales presented as five sections of the questionnaire (sections A-E), each addressing different aspects relating to SSE beliefs and sources.

**Table 1.** Details of the scales in the Irish Science Self-Efficacy Children’s Questionnaire (IS-SEC-Q) tested in Study 1.

| Scale name | Section | No. items | Assess participant's... | Origin | Changes made | Example of instructions & item |
| --- | --- | --- | --- | --- | --- | --- |
| General Academic Self-Efficacy | A | 4 | ...self-efficacy in science, literacy & maths | (20) | Split item 'Learn literacy' to 'Writing' and 'Reading' | How well can you 'Learn Science' |
| Performance-Specific SSE | B | 3 | ...self-efficacy in their academic performance in science | (20) | Changed 'science test' to 'report card' | How confident are you that you could...'Get a 5 out of 5 in science in your next report card' |
| Knowledge-specific SSE | C | 10 | ...overall self-efficacy in science knowledge. Approximates a mean score. | (20) | Changed to suit learning outcomes of Irish primary science curriculum | How well do you think you could answer a question about...'Insects & minibeasts' |
| Sources of SSE | D | 10 | ...engagement with the four sources of SSE. Approximates a mean score. Contains four subscales. | (29) | 10 items selected from original 24. Words changed from 'mathematics' to 'science' where relevant | How much do you agree with... 'Adults in my family think I'm good at learning science' |
| Task-specific SSE | E | 7 | ...overall self-efficacy to complete science tasks successfully. Approximates a mean score. | (20) | Changed to suit learning outcomes of Irish primary science curriculum (35) | How well do you think you could... 'Design an experiment to test an idea' |

SSE=Science Self-Efficacy. See S1 Figure in Supplementary Information for the entire questionnaire

In general, these questionnaires contain a series of questions, henceforth termed ‘items’. Items assessing the same latent variable are grouped together as ‘scales’. Scales that are assessing more than one latent variable are often divided into ‘subscales’. For each item, participants are asked to choose their response using one value from a provided response scale. Response scales can range from 1-3, 1-5, 1-7, 1-10 or 1-100. For the IS-SEC-Q, a 7-point response scale was chosen to a) allow respondents to make fine-grained differences between question items, b) reduce the risk of participants over-interpreting the distances between points, as is the risk with larger scales; and c) give participants a ‘neutral’ midpoint to improve scale quality (33). Each response point was uniquely labelled to increase children’s comprehension (34).

The initial IS-SEC-Q was reviewed by an expert panel of two educational researchers, one scientist facilitator and two primary school teachers (see A Fig 1), which recommended splitting item 3 (‘Learn reading, writing & literacy skills’) in the General Academic Self-Efficacy scale into two separate items: ‘Learn writing’ and ‘Learn reading’, to improve child comprehension. The items were checked for readability using an online Flesch-Kincaid test, indicating an age level of 9.4, which was appropriate for the intended sample of 11-12 years old. The questionnaire was then pre-tested with four children (three girls, aged 12 and one boy, aged 11). As all children reported to understand the questionnaire, no subsequent changes were made and the questionnaire was ready to be piloted (B Fig 1).

### Study 1: Examining the Validity of the Initial IS-SEC-Q

#### Participants & procedure

The IS-SEC-Q was completed in a paper-based format by 92 participants (52 boys and 40 girls, Mage=11.54, SD=0.58) from three different mixed-sex primary schools (one Gaelscoil, two English-speaking). Data was anonymous as participants were assigned an identifier code. Questionnaire instructions were given via Powerpoint slides.

Four participants (2 boys and 2 girls, 12 years old) who completed the IS-SEC-Q were randomly selected to participate in semi-structured interviews. Interviews were audio-recorded (length: 8-18 min) and took place on school grounds. Pseudonyms were used to assure the participants of anonymity. The word ‘confidence’ was used as a proxy for self-efficacy, as it would be more easily understood by young children (27). Participants were asked about their interpretation, comprehension and use of the questionnaire. The general interview schedule used was as follows:

1. Tell me a little bit about science at school
2. We’re going to have a look at the questionnaire you completed now.

**(Examples of probing questions):**

a. The first question asked you how well you could learn about science, maths and reading. You picked ‘very well’ for science and ‘perfectly’ for the others, do you want to tell me a bit about that?
b. It said here ‘how confident would you be if you got a 5 out of 5 in science, and you said ‘very confident’. Then why did you say ‘extremely confident’ for getting a 3?
c. For the question ‘How confident are you that you could answer questions correctly about Magnets’, you picked ‘neither poorly nor well’, do you want to tell me a little more about that?

### Data analysis

Questionnaire answers were input directly into the statistical software programme IBM SPSS Statistics version 24. The descriptive statistics of each scale was examined.

Exploratory factor analysis was performed to examine the construct validity of scales C, D and E. Construct-validity concerns the relationship between different variables within a scale. Different items assessing the same latent variable should have a good correlation with that variable, or ‘factor’, and poorer correlation with the other variables (32). The strength of this correlation is indicated by ‘factor loadings’ which range from 0-1, 1 indicating perfect correlation. Principal axis factoring was used due to its insensitivity to multivariate normality violations (36). An oblique rotation was used a) to allow for correlation between the resulting factors to minimise loss of valuable information (36) and b) because it was the rotation employed by Usher & Pajares (29) for their Sources of Self-Efficacy in Mathematics scale, which would allow for comparison of results. Factor loadings under |0.35| were suppressed as done by Usher & Pajares (29).

Once the construct validity of these scales was assessed, Cronbach’s coefficient alpha for any latent variables was calculated. Cronbach’s coefficient alpha is the most common measure of internal consistency (sometimes referred to as internal reliability) and assesses how well the items within a scale correlate with each other (37). It ranges from 0 (no correlation) to 1.00 (perfect correlation). A Cronbach’s coefficient alpha of .67-.80 is often deemed adequate in social sciences (37). It was determined that a value greater than .70 would be considered satisfactory for this study.

The construct validity of scales A or B was not examined, as the items in these scales are interpreted independently and do not represent a single latent variable.

Interviews (N=4) were transcribed verbatim. Transcripts were examined to identify any reported problems in item comprehension or interpreting/using the scale.

## Results

### Comprehension & Interpretation of the IS-SEC-Q

Analysis of the interviews did not reveal any common misinterpretations for sections A, C, D or E in the questionnaire (S1 Figure), however there was some confusion about the items in section B. Section B (the performance-related SSE scale, section B, S1 Figure) aims to assess participant’s confidence in achieving specific grades in science in their next home report cards. If answered correctly, participant’s reported confidence (*i.e.* SSE) should increase as the grade/achievement level decreases. However, the cohort mean scores indicated that participants were most confident they would achieve 4 out of 5 in science (M=5.1, SD=1.07), less confident they would achieve 3 out of 5 (M=5.01, SD=1.54) and least confident that they would achieve the top grade of 5 out of 5 (M=4.6, SD=1.10). This suggested that participants did not interpret the item instructions as intended.

This was further investigated by looking through all the questionnaire responses for the items in this section (Section B, S1 Figure). to identify different response patterns. Once the different response patterns were identified, they were quantified frequencies and percentages (Table 2). As shown in Table 2, only 30.43% of participants answered the questionnaire as expected (pattern #1, confidence for ‘Get a 3’ in science greater than ‘Get a 4’, which is greater than ‘Get a 5’). An additional 20.65% of participants reported that they were most confident about achieving a 3 out of 5 (Table 2, pattern #2), as expected, but then chose the same response on the scale for two of the items (*e.g.* the response for 3 is equal to that for 4, or 4 is equal to 5). The remaining 48.92% of participants answered this section in six unexpected ways. The largest portion of this were participants who only reported their confidence for one grade and left the other two items blank (10.87%, pattern #5 Table 2), and participants who reported that they were most confident in getting a 4 out of 5 (20.65%, pattern #3 Table 2). This suggests that these participants were not indicating their confidence in achieving each grade, but rating which grade they think that they would achieve.

**Table 2.** Response patterns in the performance-related SSE scale (section B) in Study 1.

| # | Response Patterns | N | % of participants |
| --- | --- | --- | --- |
| 1 | <sup>a</sup> Confidence for Get a 3 higher than 4, which is higher than 5 | 28 | 30.43 |
| 2 | Confidence for Get a 3 the highest, but 2 items have the same response | 19 | 20.65 |
| 3 | Confidence for Get a 4 is the highest | 19 | 20.65 |
| 4 | Confidence for Get a 3 is the same as 4 and 5 | 6 | 6.52 |
| 5 | Confidence for only 1 grade given, two are left blank | 10 | 10.87 |
| 6 | Confidence for Get a 3 is lower than 4, which is lower than 5 | 1 | 1.09 |
| 7 | Confidence for Get a 3 the lowest, but 2 items have the same response | 7 | 7.61 |
| 8 | Confidence for Get a 4 is the lowest | 2 | 2.17 |
| **Total** |  | **92** | **100** |

SSE=science self-efficacy. Response patterns indicate the ranking of the three items in the scale by participants (section B, S1 Figure). <sup>a</sup> The expected response pattern for this section in the questionnaire.

Subsequent probes about these items in the interviews (N=4) revealed that there was confusion on how to answer this scale. For example, ‘Kate’ (12 year-old girl) said “…here we got really confused about that ‘cause… I’d be very doubtful I’d get a 3 but I know I’d get higher so everyone got confused”. This further indicates that participants did not report their confidence in achieving each grade, but instead chose the highest response on the scale for the grade level that would most likely achieve. To improve participant interpretation of this scale, instructions for this section in the questionnaire were changed from ‘How confident are you that you could get these marks (out of 5) in your next report card’ to ‘How confident are you that you could get at least these marks (out of 5) in your next report card?’. In addition, the items were rephrased from ‘Get a 3 out of 5’ to ‘Get <u>at least</u> 3 out of 5’ (revised section B in S2 Figure). Specific instructions in completing this section were included in the Powerpoint presentation given to participants before completing the questionnaire

### Construct validity-Exploratory Factor Analysis

The construct validity of the Knowledge-specific SSE scale, the Task-specific SSE scale and the Sources of SSE scale was investigated using exploratory factor analysis. The results of the factor analysis for each scale are discussed in turn below (sections C, E and D, S1 Figure).

The Knowledge-specific SSE scale (section C, S1 Figure) aims to assess student’s SSE in science-related knowledge. Scale items (n=10) correspond to learning outcomes from the primary science curriculum. As the curriculum is split into four units referred to as ‘strands’ (‘Living Things’, ‘Energy and Forces’, ‘Materials’, ‘Environmental Awareness & Care’), a four-factor solution was expected. The Kaiser-Melkin-Olkin (KMO) value was greater than 0.50 (KMO=0.67), and the Bartlett’s test of sphericity was significant with p<.001 (χ2=189.28, df=45), which confirmed the sample size to be appropriate for factor analysis (38). A four-factor solution was extracted, accounting for 69% of the variance in responses. Four items loaded on Factor 1, three items loaded on Factor 2, two items loaded on Factor 3 and one item loaded on Factor 4. Factor loadings ranges for each were 0.80, 0.59-0.72, 0.59-0.80 and respectively. One item loaded on both Factor 1 and 3 (0.38 and 0.35 respectively).

Although four factors were extracted as expected, the items loading on these factors did not match the four primary science curriculum strands. To improve construct validity of this scale, three additional learning outcomes of the primary science curriculum were added as scale items to this section. To more closely reflect the curriculum (38), two items: ‘Magnets’ and ‘Electricity, batteries, bulbs & switches’ were also combined (revised section C, S2 Figure).

The Task-specific SSE scale (section E, S1 Figure) aims to assess student’s mean SSE in completing scientific tasks. As all 7 items originate from the same skill set across the ‘working scientifically’ section of the Irish primary science curriculum (35), the exploratory factor analysis was expected to reveal a one-factor solution. The KMO was 0.67 and Bartlett’s test of sphericity significant with p<.001 (χ2=215.18, df=21), again indicating an appropriate sample size for factor analysis. Exploratory factor analysis revealed a two-factor structure (64% of the total variance), which was not as expected. Five items loaded on Factor 1 and two items loaded on Factor 2. Factor loading ranges were 0.47-0.82 and 0.79-0.99 respectively. The Cronbach’s alpha coefficient for these two factors was 0.86 and 0.80 respectively. To attempt to achieve a unidimensional structure (*i.e.* all items reflect one latent variable: science skills), additional learning outcomes from the ‘Working scientifically’ section of the primary science curriculum (38) were added as scale items (revised Section E in S2 Figure).

The Sources of SSE scale (Section D, S1 Figure) aims to assess participant’s engagement with the four sources of SSE. Based on previous research (17), an exploratory factor analysis should reveal a four-factor solution, with one factor representing each SSE source. However, it revealed a two-factor solution, with one item, ‘Seeing adults do well in science pushes me to do better’ loading onto two factors. The items from Mastery Experience, Verbal Persuasion and Emotional State loaded onto Factor 1, (48% of the variance). Factor loadings ranged from 0.35-0.92. The items for Vicarious Experience loaded on Factor 2 (15% of the variance). Factor loadings ranged from 0.49-0.81. The Cronbach’s alpha value calculated for these two factors were 0.87 and 0.81 respectively, which indicate good internal consistency (>0.70).

To improve the construct validity of this scale, and potentially obtain the same four-factor solution as obtained by researchers in other contexts, 12 items from the Usher & Pajares (29) scale that were originally omitted were re-added to the revised questionnaire (items D1-16, D21-D26 in Section D, S2 Figure). In addition, to include scientists as possible providers of SSE information, four novel scientist-specific items were added: two items assessing Vicarious Experience From Scientists (D17 and D18, S2 Figure) and two assessing Verbal Persuasion From Scientists (D19 and D20, S2 Figure). These were designed by adapting items relating to adults, teachers of peers, to refer to scientists *e.g.* rewriting ‘Other students have told me that I am good at learning science’ to ‘<u>Scientists</u> have told me that I am good at learning science’.

### Conclusions from Study 1: the need to revise and re-test the IS-SEC-Q

The results from Study 1 illustrated that overall, participants understood the items in the questionnaire and children did not report any difficulty in using the 7-point Likert-like scale correctly. However, there was some confusion about how to answer the items in the performance-related SSE scale (section B in S1 Figure). Study 1 results also indicated some issues relating to the construct validity of sections C, D and E which led to various revisions to the questionnaire. Section B was re-phrased and additional instructions were included in the Powerpoint presentation, and sections C, D and E were also revised. This revised questionnaire (S2 Figure) was re-tested in Study 2.

### Study 2: Examining the Validity of the Revised IS-SEC-Q

The aim of Study 2 was to determine whether the changes made to the IS-SEC-Q improved the validity of the questionnaire (see C in Fig 1 for developmental strategy). More interviews were conducted in Study 2 than in Study 1 to take a closer look at the interpretation and comprehension of the questionnaire. In addition to re-examining the construct validity of the scales in sections C, D and E, the convergent- and criterion-related validity of these scales were also examined.

### Participants & procedures

A sample of 282 participants (138 boys, 143 girls and 1 unreported, Mage=11.79, SD=0.57) from 12 different primary schools (4 Gaelscoils, 8 English-instructed, 8 mixed-sex, 2 all-boys and 2 all-girls schools) completed the revised version of the IS-SEC-Q (S2 Figure) in the second study. Twenty-five randomly selected participants (14 girls and 11 boys, Mage=11.76, SD=0.57) completed interviews. The questionnaires were administered as in Study 1, with additional instructions for the completion of section B. Interviews were conducted as in Study 1.

### Data analysis

In Study 2, a more detailed examination of participant’s comprehension was conducted than in Study 1. Interviews were initially coded using a deductive approach. An *a priori* coding frame was established, drawing from literature surrounding questionnaire development for children and the interviews from Study 1. The coding frame used (S1 Table) had three sections: questionnaire comprehension (S1 Table A), use of the Likert-like scale (S2 Table B), and comparison with the questionnaire for confidence to ‘learn science’ (S1 Table C). First level coding was performed using NVivo and the second level coding used the ‘Comment’ feature in Word.

As in Study 1, questionnaire answers were input directly into the statistical software programme IBM SPSS Statistics version 24. Item 3 in the Mastery Experience’ subscale of the ‘Sources of Science Self-Efficacy’ scale, was reverse coded when input into SPSS as it was the only negatively phrased item in the scale (from ‘Even when I study hard I do poorly in science’ to ‘When I study hard I do well in science’). The descriptive statistics of each scale was examined.

Confirmatory factor analysis was performed with the statistical software environment R using the Lavaan package. This analysis was to determine whether the Sources of SSE scale (section D in S2 Figure) had a four-factor structure. To perform confirmatory analysis, it must be justified (either empirically or theoretically) that the structure (a.k.a. the model) being tested is likely to be present. As Usher & Pajares (29) had previously demonstrated that this scale had a four-factor structure in mathematics, this indicated that confirmatory factor analysis was an appropriate method to use. However, this analysis was done excluding the four scientist-specific items (items D17-D20 in section D, S2 Figure), as there was not enough empirical evidence prior to this study to suggest that these items would load on any of the four factors. Missing data was handled by listwise deletion (resulting in 244 complete cases), achieving sufficient Trucker’s congruence coefficient of 0.92 for factor loadings showing ‘wide’ communalities (39). Goodness of fit indices examined were the chi-square statistic, the Comparative Fit Index, the Tucker Lewis Index, the Root Mean Square Error of Approximation and the Standardized Root Mean Square Residual.

Once the confirmatory factory analysis confirmed the presence of a four-factor structure in section D (S2 Figure), an additional exploratory factor analysis was performed to investigate how the scientist-specific items (items D17-D20 in section D, Figure S2) fit into the structure. The same settings for the exploratory factor analysis were used as in Study 1. To investigate the internal consistency of the scales, Cronbach’s alpha coefficient was calculated for sections C, D and E (S2 Figure).

In addition to construct validity and internal consistency, in Study 2 the convergent-related validity and criterion-related validity of the questionnaire (S2 Figure) was also examined. Convergent-related validity is a subtype of construct-related validity (32). Similar constructs with good convergent-related validity should correlate well with each other and poorly with others. For example, the item ‘Learn science’ from section A and ‘Get a 5 out of 5’ in section B should correlate well with the total mean scores from sections C and E, and the subscales in section D. Similarly, as sections C, D and E all represent constructs linked to science self-efficacy, they should all correlate well with each other. Convergent-related validity was assessed by examining Pearson’s correlation of each participant’s scale mean with the other scale means in the questionnaire for that participant.

Criterion-related validity concerns the correlation between a measure and a ‘gold-standard’, i.e. a valid and reliable external measure of the construct of interest (32). As such a standard does not exist for science self-efficacy, to provide evidence for this a comparative analysis between the questionnaire and interviews was done. Cohen’s kappa coefficient was calculated as measure of inter-rater reliability between participant’s answers to the interview question ‘Are you confident in science?’ and their questionnaire answers to the item ‘learn science’ in section A (Section A S2 Figure). Interview responses were coded as indicating ‘High confidence’, ‘Medium confidence’ or ‘Low confidence’ (S1 Table C). Questionnaire responses were grouped into the similar categories: response options Terribly’ and ‘Very Poorly’ were grouped as ‘low confidence’, response options ‘Poorly’, ‘Neither Poorly or Well’ and ‘Well’ were grouped as ‘medium confidence’, and response options Very well’ and ‘Perfectly’ were grouped as ‘high confidence’.

## Results

### Interpretation & comprehension: the word ‘confidence’ was a good proxy for ‘self-efficacy’

To determine whether participants had a good understanding of the word ‘confidence’, all 25 interview participants were asked ‘What does confidence mean to you?’. The coding of their responses revealed two main themes. The most common theme was ‘Knowing you can do it’ (n=14): “It means that I’m able, that I’m confident that I’m able to do it, that I know that I’m able to do it….I feel good when I’ve finished and I think I’ve done a good job” (‘Eimear’, 12 year-old girl).

The second theme was ‘Feeling comfortable doing it’ (n=8): “…like being confident is when you like think you’re at something and em you’re very confident to do it so you’re not like ashamed or afraid to do it” (‘Thomas’, 12 year-old boy). As self-efficacy is linked with how you feel during the task i.e. Emotional State, this reflects ‘Knowing you can do it’ and ‘Feeling comfortable doing it’. This supports that the word confidence served well as a proxy for self-efficacy.

### Interpretation & comprehension: participants could correctly use the 7-point Likert-like scale

To confirm that participants could correctly use the 7-point response scale, they were asked a) whether they could provide a justification for their answers b) why they chose the neutral midpoint option and c) whether they could make fine-grade distinctions *i.e.* know why they chose ‘Agree’ over ‘Slightly Agree’ (S1 Table C).

- Most participants (n=24) could provide a rationale for their chosen response points on the scale.
- Most participants (n=22) chose the middle option to convey middle/average response.
- Ten participants were asked about differentiating between two scale points on similar items. Most (n=7) could provide a rationale for their choice.

These findings supported that most children interpreted the 7-point Likert-like scale correctly and were able to use it to express their opinion about each item of the questionnaire.

### Interpretation & comprehension: interpretation of the performance-related SSE scale was improved

Study 1 revealed that many participants misinterpreted the items in the performance-related scale (section B) and as a result changes were made both to the wording of the items and the accompanying administration instructions. Study 2 (N=282) saw an improvement in the user’s interpretation. As can be seen in Table 3, 50.35% of participants (an increase of 19.92% from Study 1) reported the expected pattern (pattern #1, confidence for Get a 3 higher than 4, which is higher than 5), again with some participants (26.24%) reporting pattern #2 (Confidence for Get a 3 the highest, but 2 items have the same response), similarly to Study 1 (Table 2).

**Table 3.** Response patterns for the performance-related SSE scale (section B) in Study 2. SSE=Science Self-Efficacy. Response patterns indicate ranking of items by participants. Patterns in grey were those observed for interview participants (N=25).^a^ The expected response pattern for this section in the questionnaire (S2 Figure)

| # | Response patterns | Questionnaires<br>(N=282) |  | Interviews<br>(N=25) |  | Reasoning given in<br>interviews |
| --- | --- | --- | --- | --- | --- | --- |
|  |  | N | % | N | % |  |
| 1 | <sup>a</sup>Confidence for Get a 3 higher than 4, which is higher than 5 | 142 | 50.35 | 15 | 60 | Completed as expected: not asked about reasoning in interviews |
| 2 | Confidence for Get a 3 the highest, but 2 items have the same response | 74 | 26.24 | 7 | 28 | Chose increasing confidence in decreasing grade difficult, but confidence for two grades equal |
| 3 | Confidence Get a 4 is the highest | 33 | 11.70 | 2 | 8 | Chose which grade they were more likely to achieve |
| 4 | Confidence for Get a 3 is the same as 4 and 5 | 11 | 3.90 | - | - |  |
| 5 | Confidence for only 1 grade give, two are left blank | 6 | 2.13 | - | - |  |
| 6 | Confidence for Get a 3 is lower than 4, which is lower than 5 | 8 | 2.84 | 1 | 4 | Chose which grade would make her feel the most confident if she obtained it |
| 7 | Confidence for Get a 3 the lowest, but 2 items have the same response | 6 | 2.13 | - | - |  |
| 8 | Confidence for Get a 4 is the lowest | 0 | 0.00 | - | - |  |
| 9 | All three items are missing an answer | 2 | 0.71 | - | -- |  |
| **Total** |  | **282** | **100** | **25** | **100** |  |

In the interviews (N=25), fifteen participants completed this scale as expected (pattern #1, Table 3) so were not asked about their rationale. However, ten interview participants did not complete this section as expected and were asked to provide their rationale. Two participants provided similar reasons to those proposed by others in Study 1: they selected the grade level which they think they would achieve (pattern #3, Table 3). Pattern 5 could have similar explanation but none of the interviewees had adopted this pattern so this cannot be confirmed.

One interview participant chose the answer according to how confident she would feel after receiving each grade i.e. she would feel the most confident in science if she received a 5 out of 5 (pattern #6, Table 3). Seven interview participants completed the scale with their confidence decreasing with increasing grade difficulty, but with two of items being equal (*e.g.* they were equally confident about achieving a 4 as they were about achieving a 5, pattern #2, Table 3): “Because I wouldn’t really think I’d get a really good science score so I’d say 4 out of 5 or 3 out of 5” (‘Jimmy’, 12 year-old boy).

This suggests that these participants viewed ‘Get a 4 out of 5’ and ‘Get a 5 out of 5’ to be of equal difficulty. This is somewhat reflected in the mean scores for this section with the difference in cohort scale means for ‘Get a 3’ being twice that of ‘Get a 4’ and ‘Get a 5’ (Table 4). Overall, most questionnaire participants completed this scale as expected (76.59% most confident about receiving a 3 out of 5 than a 5 out of 5, patterns #1 and #2, Table 3), which suggests that additional administration instructions and rewording of the scale aided in scale interpretation.

**Table 4.** Cohort means and standard deviations for the items in section B of the IS-SEC-Q (Study 2)

| Section B Scale Item | Mean | Std. Deviation |
| --- | --- | --- |
| Get <u>at least</u> 5 out of 5 | 4.45 | 1.26 |
| Get <u>at least</u> 4 out of 5 | 5.40 | 1.04 |
| Get <u>at least</u> 3 out of 5 | 5.81 | 1.40 |

Although 23.41% of questionnaire participants did not follow this pattern, justifications from interviews suggest that the reasoning for ‘Get a 5 out of 5 in science’ alone is interpreted correctly. Only this item is used in the subsequent correlation analysis assessing convergent-related validity of sections C, D and E so it should not be adversely affected.

### Construct validity: the internal structure of the Knowledge-specific SSE scale remains unclear

As in Study 1, an exploratory factor analysis was performed to assess the construct validity of section C: the Knowledge-specific scale. The analysis extracted a four-factor structure, accounting for 56% of the total variance (Table 5). However, similar to in Study 1, these four factors still did not reflect the four strands in the Irish primary science curriculum (Table 5). As any factor loadings below |0.35| were supressed, two items: ‘what happens to things when you mix them together’ and ‘having a balanced & nutritious diet’, did not present at all.

**Table 5.** Factor loadings of the Knowledge-specific Science Self-Efficacy scale (section C) from the Irish Science Self-Efficacy Children’s Questionnaire (Study 2).

| Scale items | Irish Primary Science Curriculum Strand | Factor Loadings |  |  |  |
| --- | --- | --- | --- | --- | --- |
|  |  | 1 | 2 | 3 | 4 |
| Plants & how they grow | Living Things | .42 |  |  |  |
| Different sources of energy | Energy & Forces | .66 |  |  |  |
| Different types of energy | Energy & Forces | .81 |  |  |  |
| Magnets bulbs switches & electricity | Energy & Forces | .79 |  |  |  |
| Lights mirrors & shadows | Energy & Forces | .65 |  |  |  |
| Wind and levers | Energy & Forces | .89 |  |  |  |
| Solids, liquids & gases | Materials | .35 |  |  |  |
| Natural & manufactured materials | Materials |  | .38 |  |  |
| The work of scientists in past and present | EA&C |  | .62 |  |  |
| Using science in your everyday lives | EA&C |  | .85 |  |  |
| How science has made the world better | EA&C |  | .75 |  |  |
| How the human body works | Living Things |  | .59 |  |  |
| How the body protects itself from disease | Living Things |  | .42 |  |  |
| Saving energy & recycling | EA&C |  |  | .87 |  |
| What happens to things when you heat and cool them | Materials |  |  | .51 |  |
| How to look after the environment | EA&C |  |  | .62 |  |
| Insects & minibeasts | Living Things |  |  |  | .51 |
| Animals from around the world | Living Things |  |  |  | .82 |
| What happens to things when you mix them together | Materials | - | - | - | - |
| Having a balanced & nutritious diet | Living Things | - | - | - | - |

EA&C=Environmental Awareness & Care. Knowledge-specific Science Self-Efficacy scale is presented in S2 Figure. Items originate from the strands of the Irish Primary Science Curriculum (35). Factor analysis was performed using principal axis factoring with promax rotation. Factor loadings greater than |0.35| were suppressed. Non presenting loadings are denoted by '-'

Exploratory factor analyses from both studies 1 and 2 strongly suggest that section C, as it exists in relation to the Irish primary science curriculum, is not a unidimensional construct. Further development of this scale would be needed for it to be used to assess participant’s SSE for specific strands of the curriculum.

This ‘mixing’ of items from different curriculum strands loading onto the same factors could be due to how science is taught in the Irish primary school curriculum. Science is not taught as an individual subject but is combined with Social and Environmental studies. In addition, teachers are encouraged to employ a cross-subject approach when teaching (35).

### Construct validity: Task-specific SSE scale obtained a unidimensional structure

As in Study 1, an exploratory factor analysis was performed to determine whether the Task-specific SSE scale with additional items (section E, S2 Figure) reflects the ‘working scientifically’ skills as a unidimensional construct. Exploratory factor analysis performed on section E extracted a clear one-factor solution, as expected, which was an improvement from Study 1. Factor loadings ranged from 0.57-0.77 (52% of total variance). The Cronbach’s alpha of 0.90 was also an improvement from Study 1. This indicated that this scale was ready for use.

### Construct validity: confirmatory factor analysis of Sources of SSE scale

To ascertain whether section D had a four-factor structure, as previously determined (29), a confirmatory factor analysis was performed to test a four-factor model (Fig 2). To compare with Usher & Pajares (29), this analysis excluded the four novel scientist-specific items.

**Fig 2.**
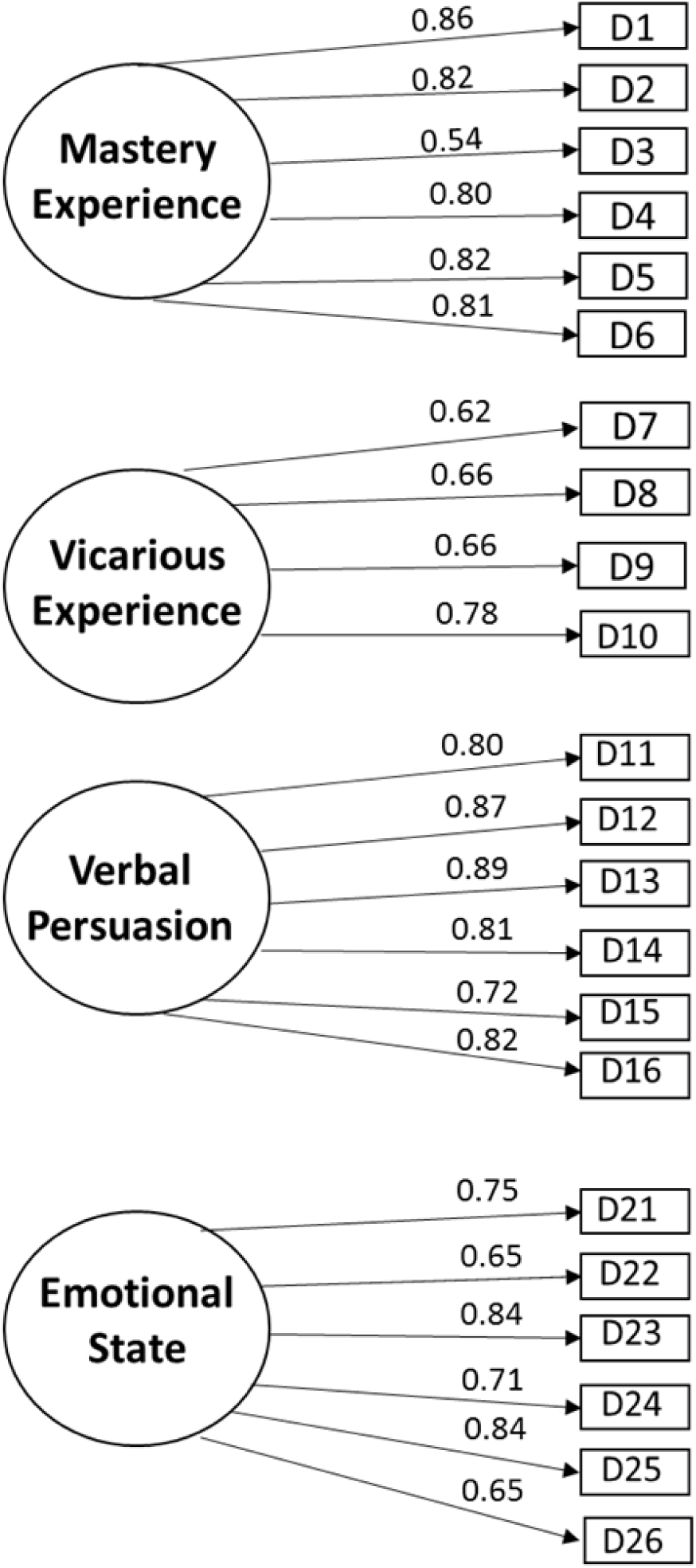
Path diagram showing factor loadings for the 22-item Sources of Science Self-Efficacy scale. An exploratory factor analysis was performed on the items in section D of the revised Irish Science Self-Efficacy Children’s Questionnaire in Study 2 (S2 Figure) to determine the internal structure of the scale. This analysis excluded the four novel scientist-specific items in order to compare it with analyses performed by Usher and Pajares (29). Circles represent latent variables. Squares represent items in the scale. Factor loadings for each item are shown above the arrows between the item and respective latent variable.

The goodness of fit indices outlined in Table 6 showed an acceptable fit for this model. Values for the goodness of fit indices are also comparable to those reported by Usher & Pajares (29). This result supports that this section contains four subscales, each representing one of the four sources of SSE, and is therefore valid in an Irish primary context.

**Table 6.** Goodness of fit indices from confirmatory factor analysis for a four-factor model for the Sources of Self-efficacy Scale

| Goodness of fit index | Cut-off values | Usher & Pajares (29) | Study 2 |
| --- | --- | --- | --- |
| Chi-square ($\chi^2$) Statistic | Ratio of $\chi^2$ to $df \leq 2$ or | 601.21 | 366.177 |
| Comparative Fit Index | $\geq .90$ for acceptance | .96 | .95 |
| Tucker Lewis Index | $\geq .90$ for acceptance | - | .94 |
| Root Mean Square Error of | $< .6$ to $.8$ | .04 | .06 |
| Standardized Root Mean | $\leq .8$ | .04 | .050<sup>a</sup> |

<sup>a</sup> Root Mean Square Residual confidence intervals are .048 and .067 with p value at $p < .05 = .098$ .

### Addition of scientist-specific items resulted in five-factor structure for the Sources of SSE scale

The four new scientist-specific items were not included in the previous confirmatory factor analysis on section D, as there was no prior empirical evidence to suggest where they would fit into the structure. To ascertain where these new items fit within section D, an exploratory factor analysis was performed with the 26 items of this section. The KMO was 0.90 and Bartlett’s test of sphericity was significant with p<.001. The exploratory factor analysis revealed a five-factor solution accounting for 67% of the total variance (Table 7).

**Table 7.** Factor loadings of the Sources of Science Self-efficacy Scale (section D).

| Sources of Science Self-Efficacy items | Extracted factors and explained variances (%) |  |  |  |  |
| --- | --- | --- | --- | --- | --- |
|  | 1 (36%) | 2 (13%) | 3 (8%) | 4 (5%) | 5(4%) |
| **Four subscales confirmed by confirmatory factor analysis** | **Factor Loadings** |  |  |  |  |
| Emotional State (D21-26) | (0.63-0.81) |  |  |  |  |
| Mastery Experience (D1-D6) |  | (0.50-0.86) |  |  |  |
| Verbal/Social Persuasion (D11-D16) |  |  |  | (0.46-0.83) |  |
| Vicarious Experience from non-scientists (D7-D10) |  |  | (0.64-0.74) |  |  |
| **Four new scientist-specific items** |  |  |  |  |  |
| D17. <sup>a</sup>Seeing scientists do well in science pushes me to do better |  |  | 0.74 |  |  |
| D18. <sup>a</sup>When I see how a scientist does an experiment I can picture myself doing the experiment in the same way |  |  | 0.70 |  |  |
| D19. <sup>a</sup>Scientists have told me that I am good at learning science |  |  |  |  | 0.98 |
| D20. <sup>a</sup>Scientists have praised me for my ability in science |  |  |  |  | 0.86 |

See section D in S2 Figure. Performed using principal axis factoring with an oblique promax rotation. Factor loadings less than |0.35| were suppressed.
Kaiser-Meyer-Olkin Measure of Sampling Adequacy=.90, Bartlett's test of sphericity significant with $p < .001$
<sup>a</sup> New scientist-specific items.

As seen in Table 7, the two items approximating Vicarious Experience From Scientists loaded on Factor 3 along with the other Vicarious Experience items (factor loadings of 0.74 and 0.70). The two items approximating Verbal Persuasion From Scientists loaded alone on Factor 5 (factor loadings of 0.98 and 0.86). This suggests that participants perceived the new items approximating Verbal Persuasion From Scientists differently to the other items assessing Verbal Persuasion.

### Interpretation & comprehension: first reports of items from the Emotional State subscale being misinterpreted

Interview responses indicated that some items from the Emotional State subscale in section D (S2 Figure) were sometimes misinterpreted by the participants. Approximately 22% (n=64) of participants (n=282) chose responses from the ‘agree’ end of the scale, reporting to experience negative emotions such as anxiety whilst doing science. Fourteen such participants were asked about their reasoning and interpretation of the items during the interview. The most commonly misinterpreted item for interview participants was the item: ‘My mind goes blank and I am unable to think clearly when I do science’. Seven of the interview participants agreed with this item to some degree because they thought it meant that they daydream during science: “…yeah I think I just get sidetracked and it kinda goes in one ear and comes out the other a lot of times” (‘Emily’, 12 year-old girl).

Another commonly misinterpreted item in the Emotional State subscale was the item ‘Doing science work takes all my energy’. Four participants interpreted it as ‘Pat’ (12-year old boy) did: “…because sometimes you do experiment where you run around the classroom for break with forces and stuff so it takes a little bit of my energy”. This seems to be the first investigation into the comprehension of these items by children, and the first report of possible misinterpretation. The percentage of participants who were queried about their interpretation of the items in the Emotional State subscale represented a fifth of participants who chose responses from the ‘agree’ end of this scale. Therefore, researchers should consider adapting or removing these items from the scale in future studies.

### Criterion-related validity

Criterion-related validity refers to how well the responses from a scale correlates to another comparable measure (32). To examine the criterion-related validity for the item ‘learn science’ in section A (S2 Figure), questionnaire responses to this item were grouped into categories (Low confidence, Medium confidence, High confidence, as described in Study 2 Data analysis) and compared with the answers from participant’s interviews to the question ‘Are you confident in science?’ (Table 8). Pupil’s responses to the question in the interviews were coded as ‘low confidence’, ‘medium confidence’ and ‘high confidence’ (S1 Table C). Cohen’s Kappa coefficient was calculated as a measure of agreement between the two responses. Cohen’s kappa coefficient was 0.40 (percentage agreement=62.5%), showing a moderate agreement between the interview and questionnaire. Although the agreement could be improved, it does give some indication that the item ‘learn science’ gives a reasonable approximation of children’s science confidence.

**Table 8.** Crosstabulation of coded answers from interview and IS-SEC-Q on confidence in science.

| Source of data | Questionnaire responses |  |  |  |  |
| --- | --- | --- | --- | --- | --- |
|  | Category | Low | Medium | High | Total |
| Interview responses | Low | 1 | 2 | 1 | 4 |
|  | Medium | 0 | 11 | 1 | 12 |
|  | High | 0 | 5 | 4 | 9 |
|  | **Total** | 1 | 18 | 6 | 25 |

Questionnaire responses are coded answers to item 'Learn Science' in section A of the IS-SEC-Q (S2 Figure). Interview responses are coded replies to the question 'are you confident in science?'.

### Convergent validity

To assess the convergent validity of the scales in the IS-SEC-Q, Pearson’s correlations were calculated between the items ‘Learn science’ (section A), ‘Get a 5 out of 5’ (section B), the mean scores of sections C and E and the subscales of Section D (Table 9). All correlations were found to be statistically significant with p<0.01. There were moderate correlations between all scales. Mastery Experience had strong correlations with ‘Learn science’, ‘Get 5 out of 5’, Knowledge-specific SSE and Task-specific SSE (r=.60, .60, .70 and .66 respectively) in agreement with other studies finding Mastery Experience to be a strong predictor of science self-efficacy (25). Inversely, Mastery Experience had a weak correlation with ‘Learn Maths’, ‘Learn Writing’, ‘Learn Reading’ and ‘Get 3 out of 3 in science’ (r=.38, .16, .31 and .29 respectively).

**Table 9.**
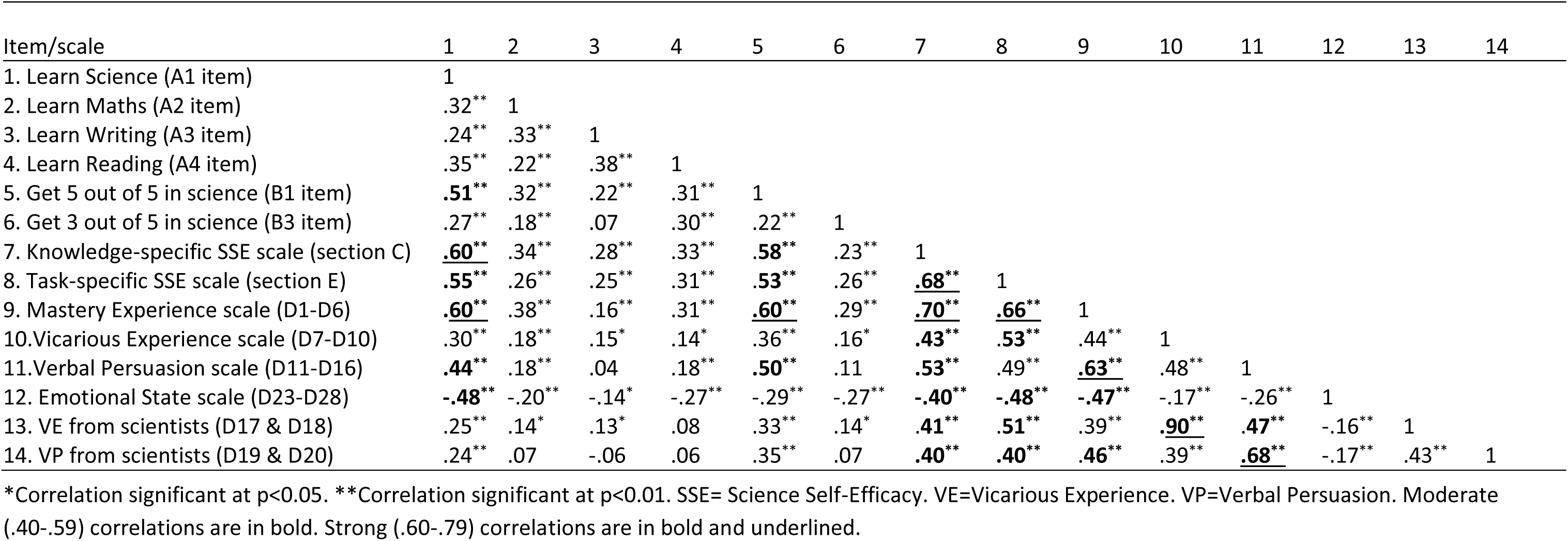
Inter-scale Pearson correlations for the Irish Science Self-Efficacy Children’s Questionnaire (study 2)

Regarding the new scientist-specific items, the Verbal Persuasion had the strongest correlation with Verbal Persuasion From Scientists (r=.68), highlighting their similarity (Table 9). Similarly, the items assessing Vicarious Experience From Scientists correlated very strongly with Vicarious Experience (r=.90). Emotional Stat’ had a negative correlation with all other subscales in agreement with previous studies (23).

The moderate-strong inter-scale correlations from this analysis supports that the different scales and subscales within the IS-SEC-Q all relate to the SSE of Irish Primary children. Furthermore, the results are in agreement with those obtained in self-efficacy studies in other contexts (20, 29), indicating that the adaptations made here have not affected the psychometric quality of the scales.

### Conclusions & Future Directions

This study aimed to develop a questionnaire, suitable for use by scientist facilitators of hands-on outreach, to assess the strengths and sources of upper primary school pupil’s science self-efficacy beliefs. The questionnaire needed to relate the Irish primary science curriculum, assess the four sources of science self-efficacy as outlined by Bandura (11), include scientists as providers of Vicarious Experience and Verbal Persuasion. To ascertain the suitability of the questionnaire as a measurement instrument of SSE beliefs, this study evaluated its construct-, convergent- and criterion-related validity. After one cycle of revisions, which were evaluated in Study 2, the resulting IS-SEC-Q presented in S2 Figure comprises 63 items divided across five sections, each answered using an individually labelled 7-point Likert-like scale. Overall, pupils could correctly comprehend and interpret the questionnaire, and use the Likert-like scale accurately to denote their responses.

Four sections of the questionnaire specifically relate to assessing the strength of pupil’s self-efficacy beliefs. The General Academic Self-Efficacy scale (Section A) assesses pupil’s self-efficacy to learn different academic subjects, to compare with their self-efficacy for science. The Performance-Related SSE assess pupil’s confidence to achieve different grades in science in their next school report. Sections C and D specifically relate to the Irish primary science curriculum. Section C (20 items) contains the Knowledge-specific SSE scale, which assess pupil’s self-efficacy to answer questions on learning outcomes from the science curriculum. Although interviews did not reveal any problems with comprehension or interpretation of these items, a clear underlying latent structure for this scale could not be obtained. This may be due to the integrative way these subjects are taught. However, the overall cohort mean for this scale does correlate well with the item ‘Learn Science’ from Section A, which suggests that if pupils are confident in answering their science questions, they are also confident in learning science in general. Whilst further work would be needed to uncover an underlying latent structure for this scale, the means for the individual items of this scale may be interpreted without caution.

Section E (10 items) contains the Task-specific SSE scale, which assess pupil’s self-efficacy to perform ‘working scientifically’ skills from the Irish primary science curriculum. As with section C, there were no issues with pupil’s comprehension or interpretation of the items in this scale. Furthermore, a unidimensional structure was obtained with exploratory factor analysis in Study 2, which indicated that all items were related to the same latent variable. The scale mean correlated well with the item ‘Learn Science’. It also correlated strongly with the mean of the Knowledge-specific scale, which is expected as they originate from the same curriculum. This scale is ready for use without any further improvement.

Section D of the questionnaire assesses the sources of pupil’s SSE beliefs, section D. This section contains an adapted version of Usher & Pajares Sources of Science Self-Efficacy scale for Mathematics (29). Confirmatory factor analysis indicated that the rewording of the items from ‘mathematics’ to ‘science’ did not change the four-factor internal structure and hence the scale is validated in the Irish primary science context. Additionally, all subscale means correlate well with the other science-related SSE items in the questionnaire, and especially well with the Knowledge- and Task-specific scales. This indicates that pupils who receive more of these sources of SSE, feel more confident in the learning outcomes of the Irish primary science curriculum.

The scale in Section D now contains new scientist-specific items, two approximating Vicarious Experience from Scientists and two approximating Verbal Persuasion from Scientists. The two items from Vicarious Experience from Scientists loaded with the other Vicarious Experience items, whilst the two Verbal Persuasion from Scientists items loaded on a new fifth factor. This suggests that children find Verbal Persuasion from Scientists to be distinctive from other non-scientist social models such as peers, teachers and family members. The means of Verbal Persuasion from Scientists correlate significantly with the other science-related means in the questionnaire, which supports its validity as a source of SSE. A potential limitation of this new latent variable is that two items may not be enough to represent a construct. Therefore, it would be advisable for future studies to add a third item assessing Verbal Persuasion from Scientists to rectify this.

Whilst the Sources of SSE scale is section D of the IS-SEC-Q has good construct- and convergent-related validity, the interviews from Study 2 revealed a potential for misinterpretation of the items in the Emotional State subscale never reported before. This should be taken in consideration when analysing these items. Future researchers focusing on Emotional State should consider to adapt, remove or further investigate participant’s understanding of these items for their studies.

Overall, the IS-SEC-Q demonstrated good psychometric properties for the assessment of the strength and sources of 11-12 years old children’s SSE beliefs, as they relate to the Irish primary science curriculum. It is now ready for use by other researchers and education practitioners to assess the SSE beliefs of upper primary school children in Ireland, either in the formal or informal science context. The IS-SEC-Q will be used in a future study to investigate the SSE of upper primary students in Ireland and to explore the effect of participation in scientist-led outreach.

## Author Contributions

**(CRediT)** Conceptualization (SC, VMC, MG), Data Curation (SC), Formal Analysis (SC), Funding Acquisition (MG) Investigation (SC), Project Administration (SC), Supervision (VMC, MG), Resources (MG), Validation (JS), Visualisation (SC), Writing (Original Draft Preparation) (SC), Writing (Review & Editing) (JS, VMC, MG).

## Ethics Statement

This work was given ethical approval by the Research Ethics Committee at the National University of Ireland Galway. The approval reference is 17-NOV-02. The Human Subjects in this research are children participants (under the age 18). Only children who return completed written positive assent forms (from themselves) and positive consent forms (from either parents/guardians) participated in the studies outlined in this manuscript. Questionnaires completed by participants were anonymous and interviews were completed using a pseudonym.

## Acknowledgements

This research was conducted in M.G’s research group within the Biochemistry discipline of School of Natural Sciences at the National University of Ireland Galway in Ireland. The research in M.G’s group is linked to the activities of Cell EXPLORERS (www.cellexplorers.com), an Educational Outreach/Public Engagement programme funded by the Science Foundation Ireland (www.sfi.ie) Discover awards 16DP4123 and 18DP5772 and materially supported by the Biochemistry discipline, the School of Natural Sciences, the College of Science and National University of Ireland Galway (www.nuigalway.ie). S.C’s PhD is funded by a College of Science scholarship supported by the Galway University Foundation (www.guf.ie). The authors thank the primary school pupils and teachers from the thirteen schools which participated in this study.

## Supporting information

### Supporting information captions

**S1 Fig. The initial Irish Science Self-Efficacy Children’s Questionnaire (IS-SEC-Q).** This is the initial IS-SEC-Q, as administered in Study 1 (4 pages long). Page 1 details the questionnaire instructions, illustrates an example and also collects covariate variables ‘Age’ and ‘Gender’. The IS-SEC-Q was designed to be printed as an A3 booklet, to maximise ease of use for the target age group.

**S2 Fig. The revised Irish Science Self-Efficacy Children’s Questionnaire (IS-SEC-Q).** This is the final IS-SEC-Q, as administered in Study 2 (7 pages long). Page 1 details the questionnaire instructions, illustrates an example and also collects covariate variables ‘Age’ and ‘Gender’. The IS-SEC-Q was designed to be printed as an A3 booklet, to maximise ease of use for the target age group.

**S1 Table 1. Coding scheme used for code interviews from Study 2. S1 Table A.** Questionnaire comprehension. **S1 Table B.** Justifying response for choosing the middle option on the Likert-like response scale**. S1 Table C.** Comparison with questionnaire for confidence to ‘learn science’

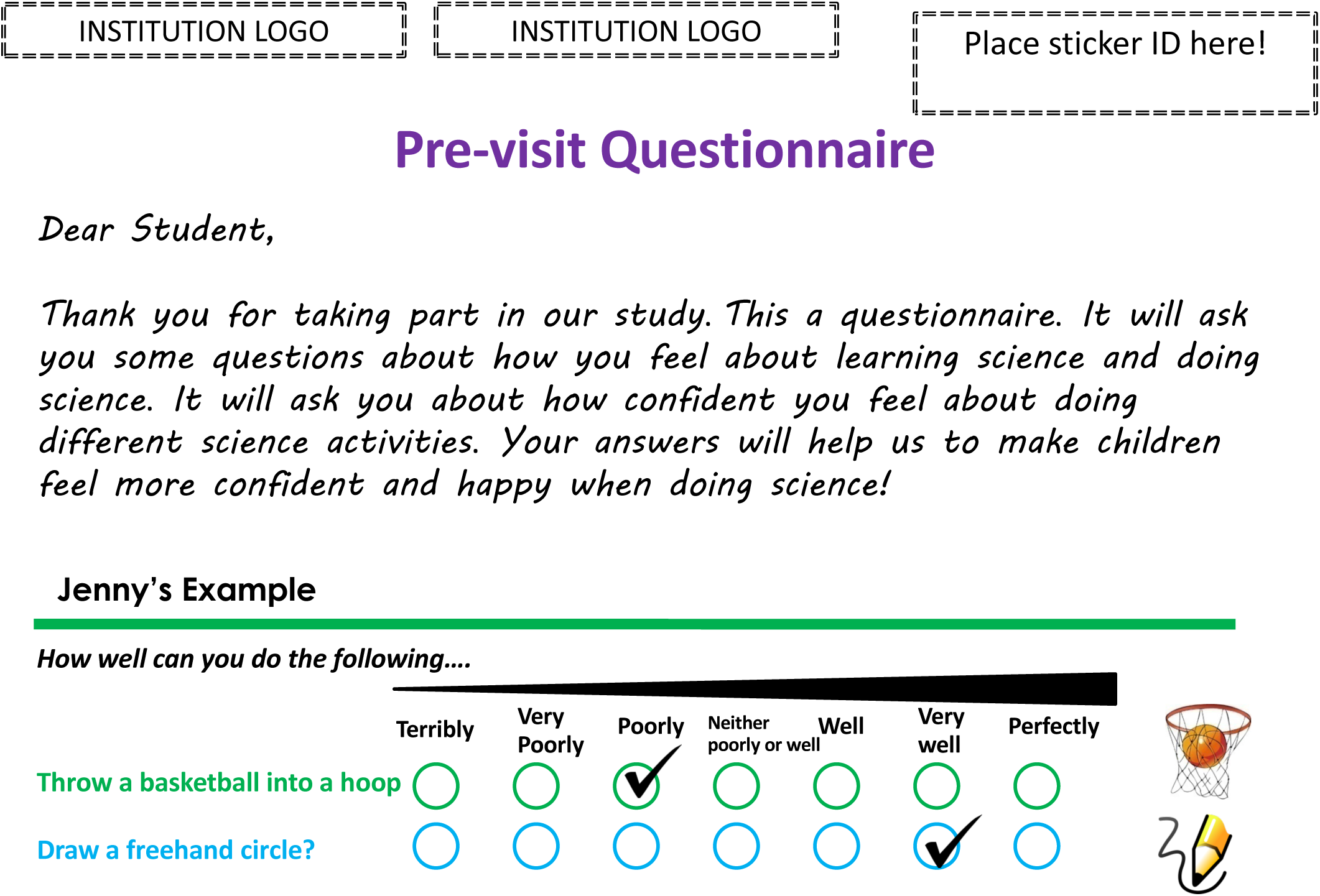

### Throw a basketball into a hoop?

Jenny is not very good at playing basketball and feels like she would miss the hoop more often than score if she tried. So she picks “Poorly” for the first question.

### Draw a freehand circle?

Jenny is really good at art and often draws pretty good freehand circles. They're not perfect though, so she picks “Very well”.

### Remember!

- Tick which circle applies to you
- Answer the following questions as honestly as you can. This is not a test!
- If you are not sure about what to do, please raise your hand for help!
- Do not copy answers from classmates, we want your answers! There are no “right” or “wrong” answers!
- If there is a question that you do not want to answer-no problem! Just skip it!

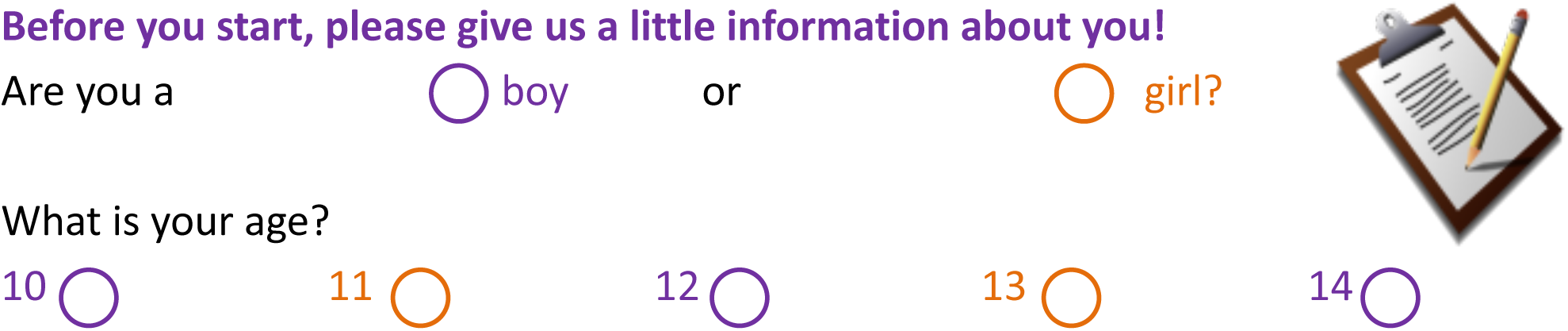
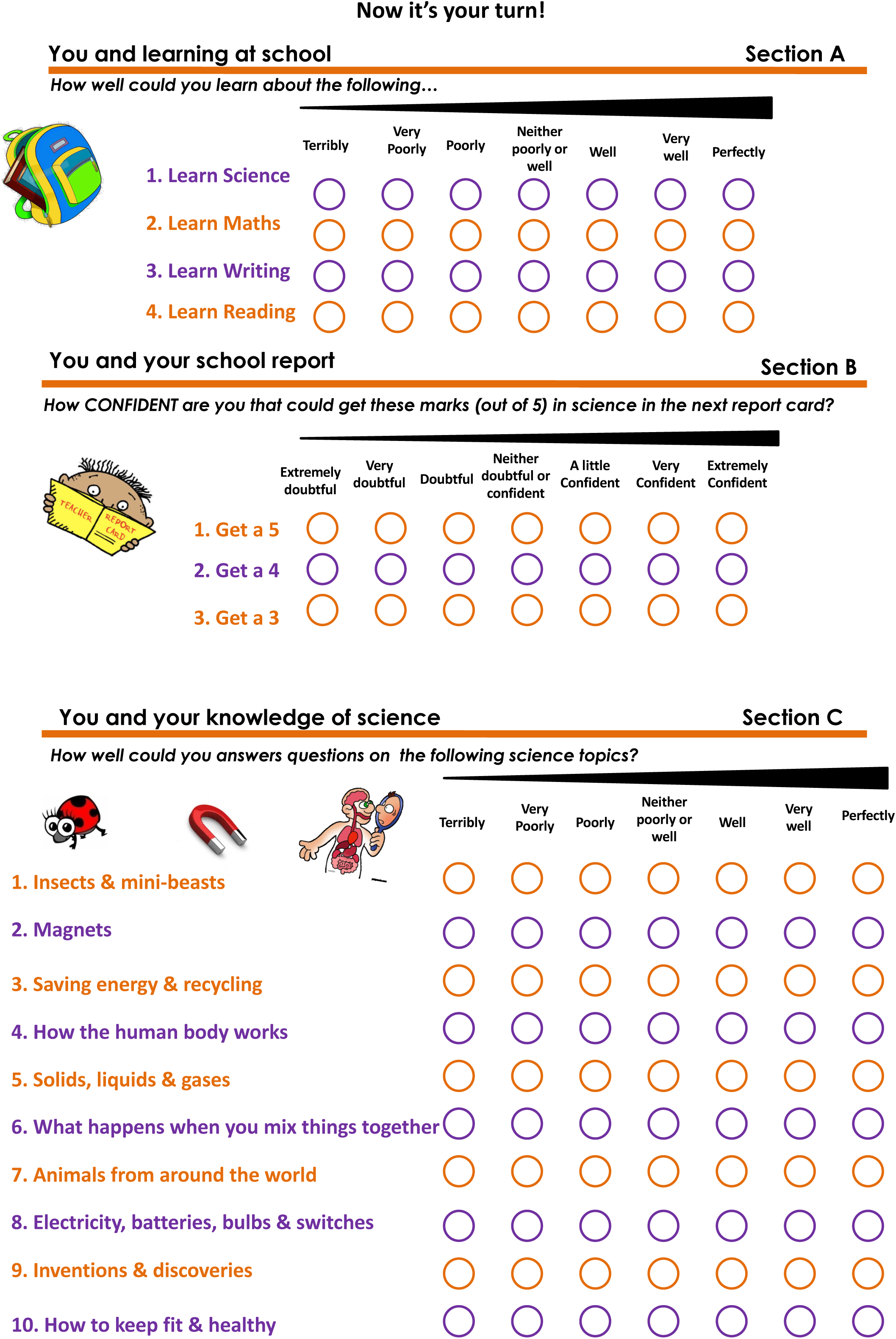
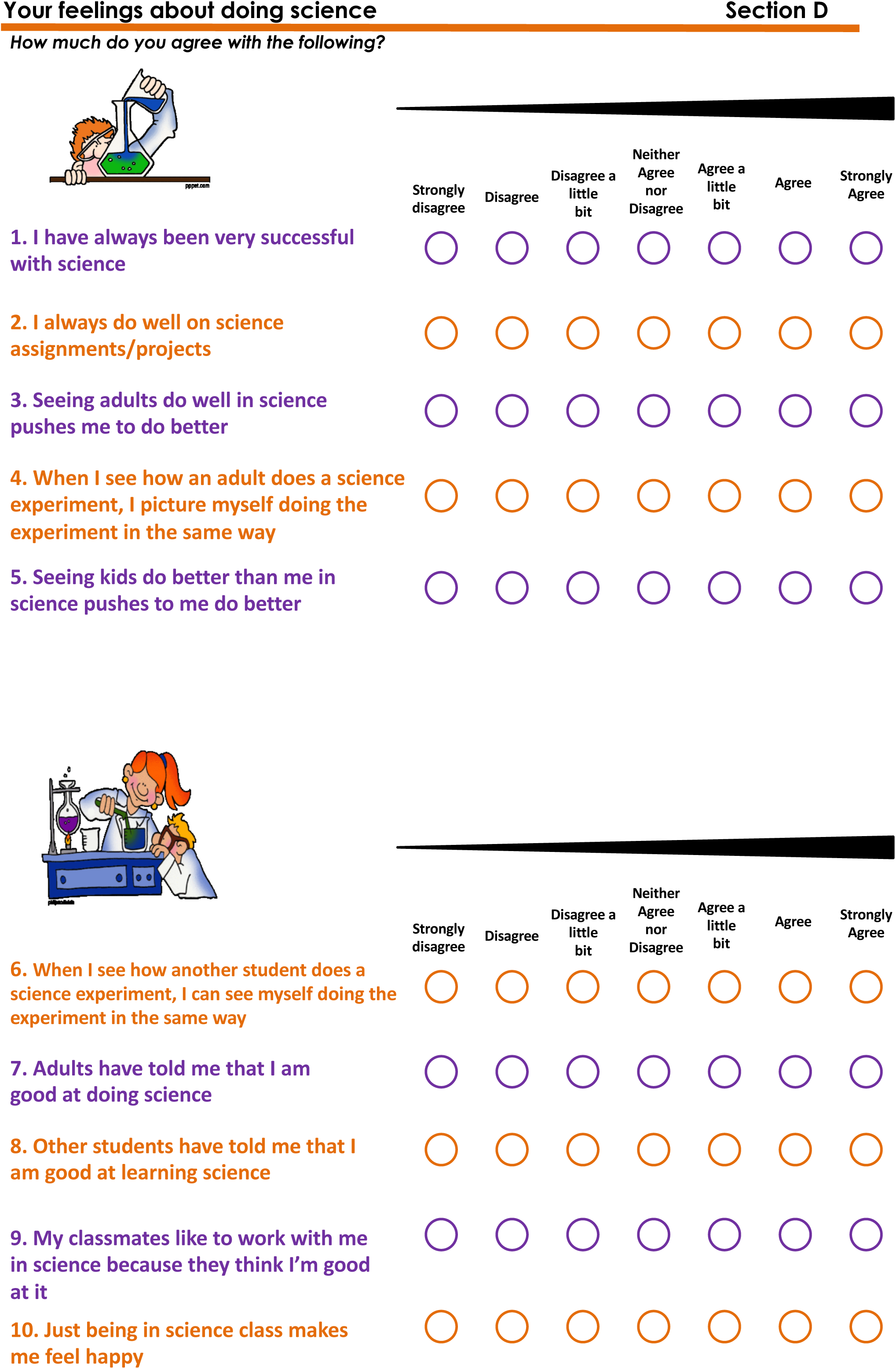
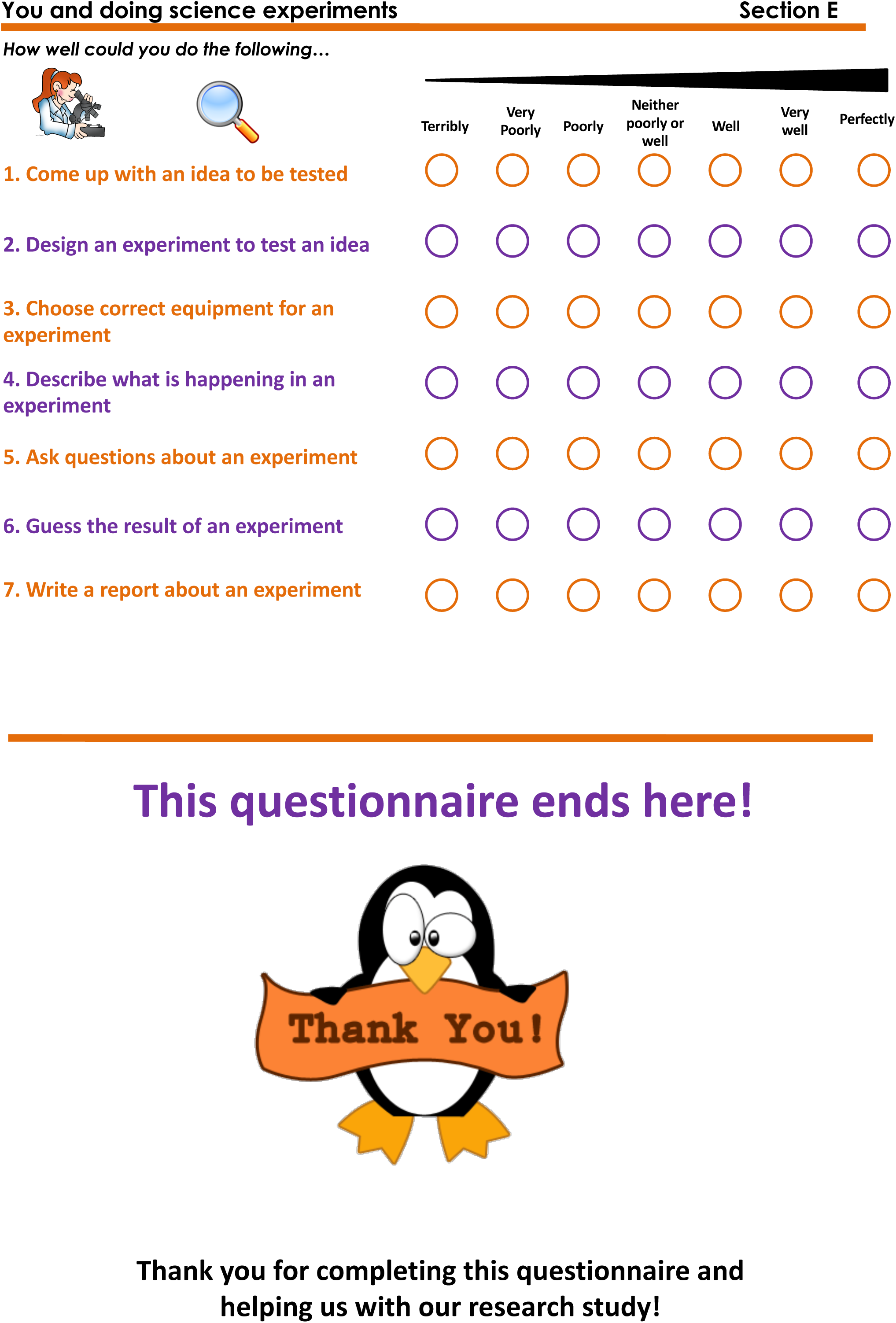

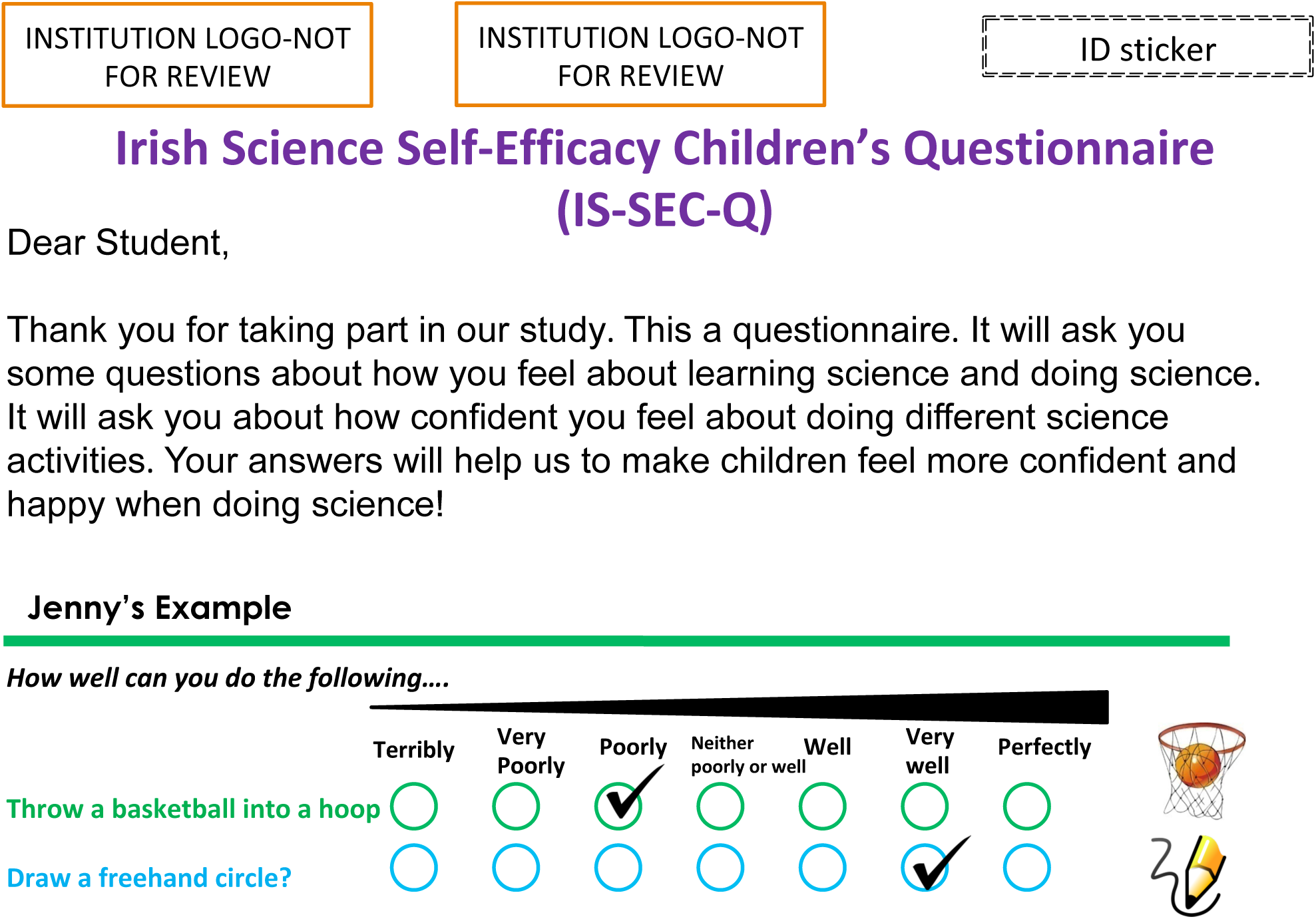

### Throw a basketball into a hoop?

Jenny is not very good at playing basketball and feels like she would miss the hoop more often than score if she tried. So she picks “Poorly” for the first question.

### Draw a freehand circle?

Jenny is really good at art and often draws pretty good freehand circles. They're not perfect though, so she picks “Very well”.

### Remember!

- Tick which circle applies to you
- Answer the following questions as honestly as you can. This is not a test!
- If you are not sure about what to do, please raise your hand for help!
- Do not copy answers from classmates, we want your answers! There are no “right” or “wrong” answers!
- If there is a question that you do not want to answer-no problem! Just skip it!

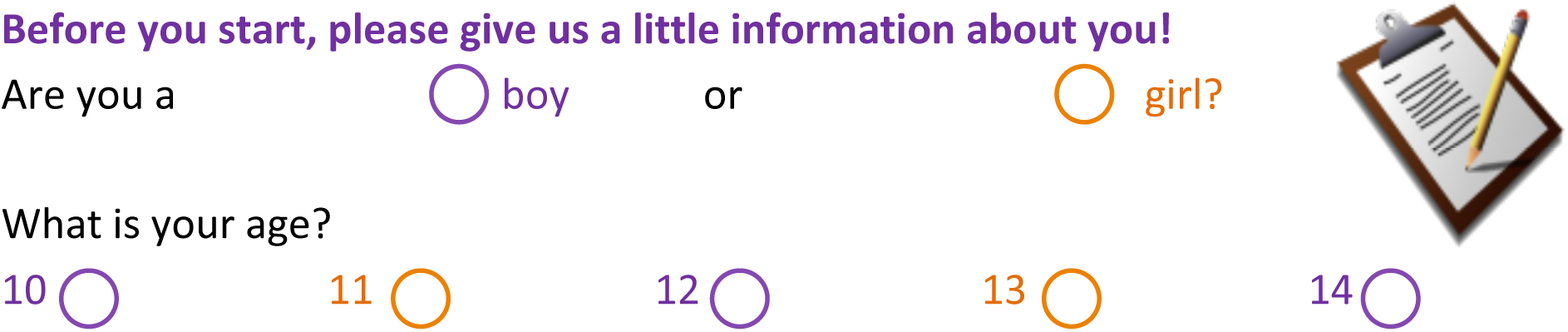
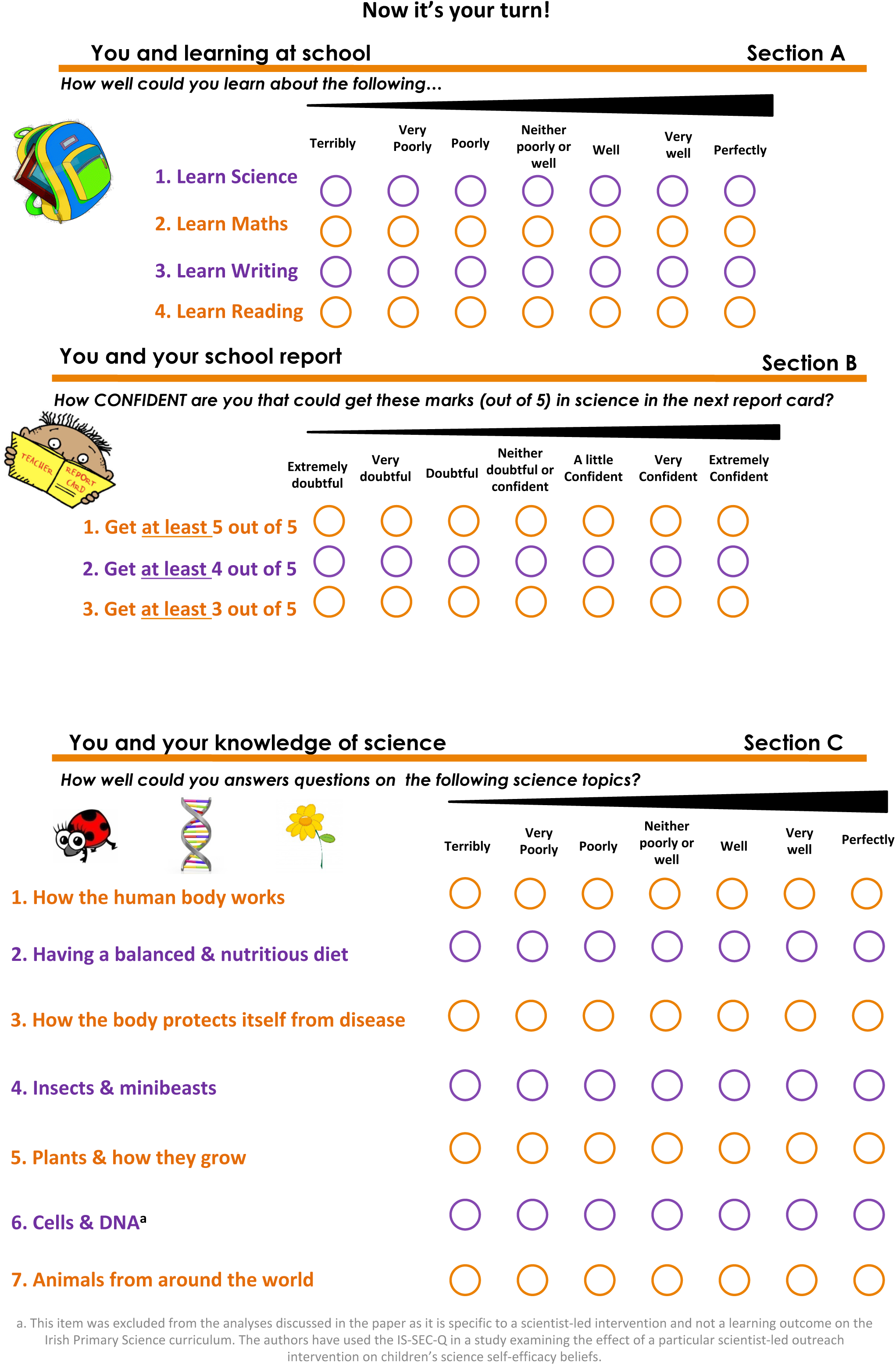
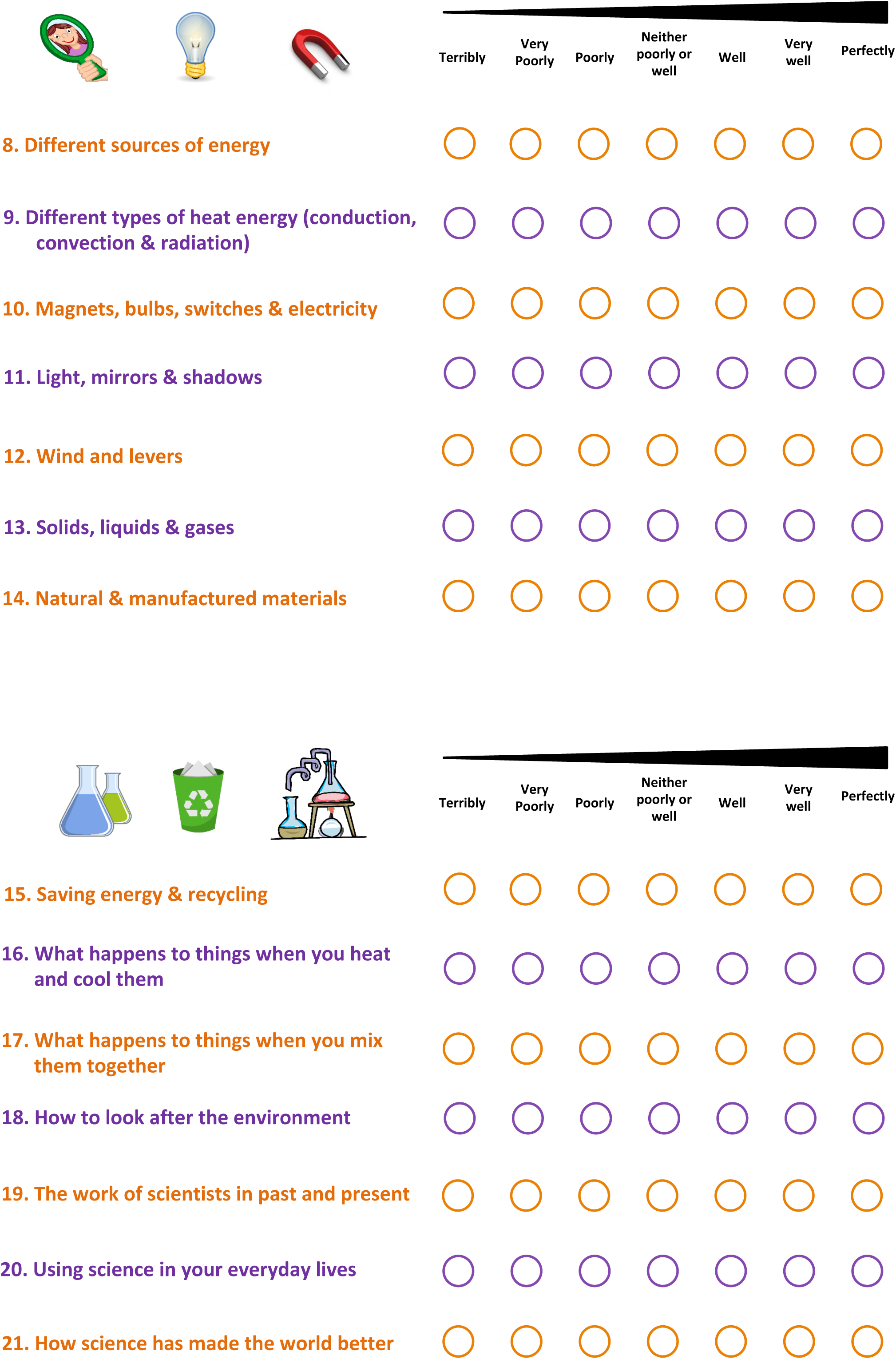
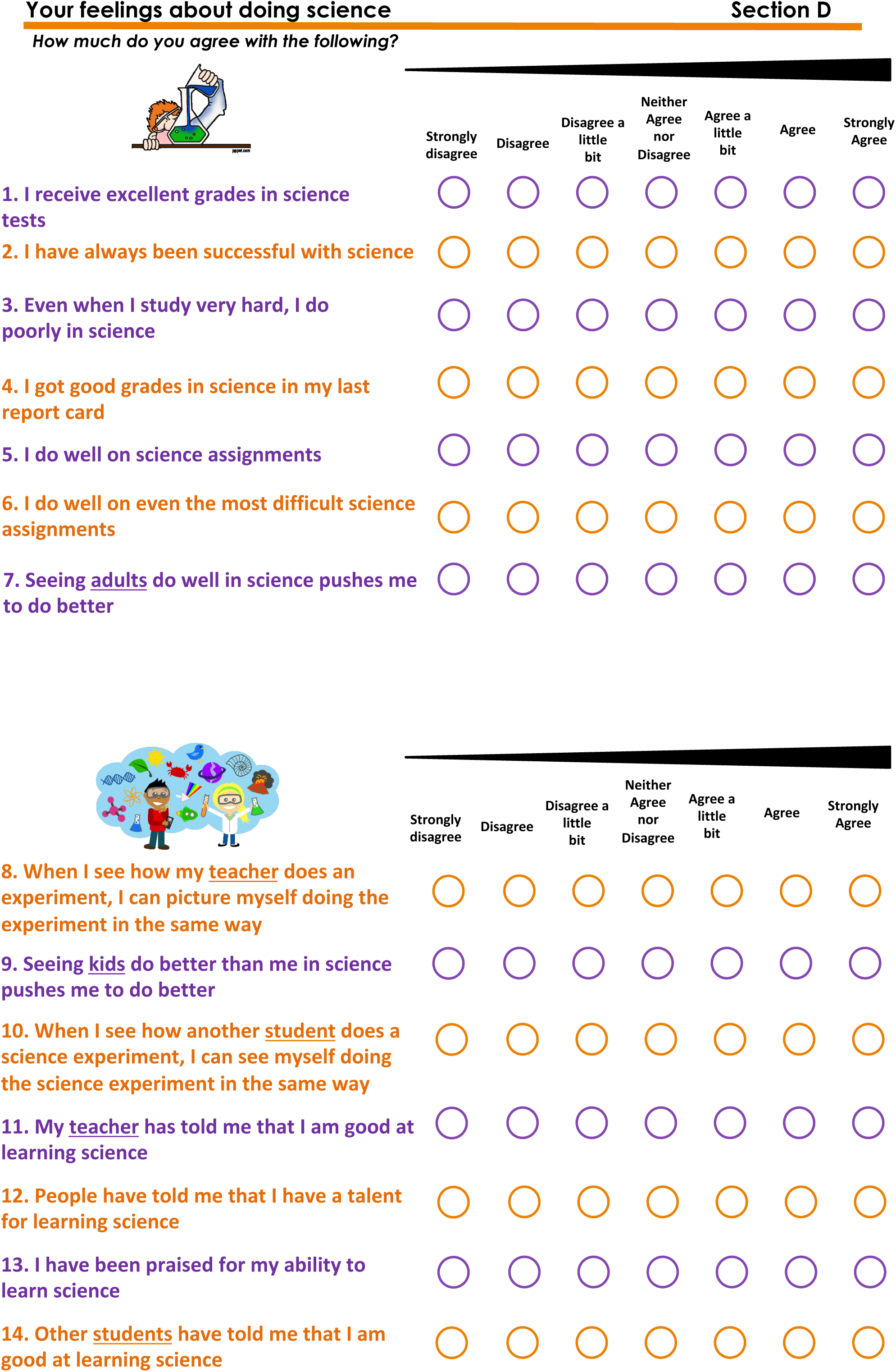
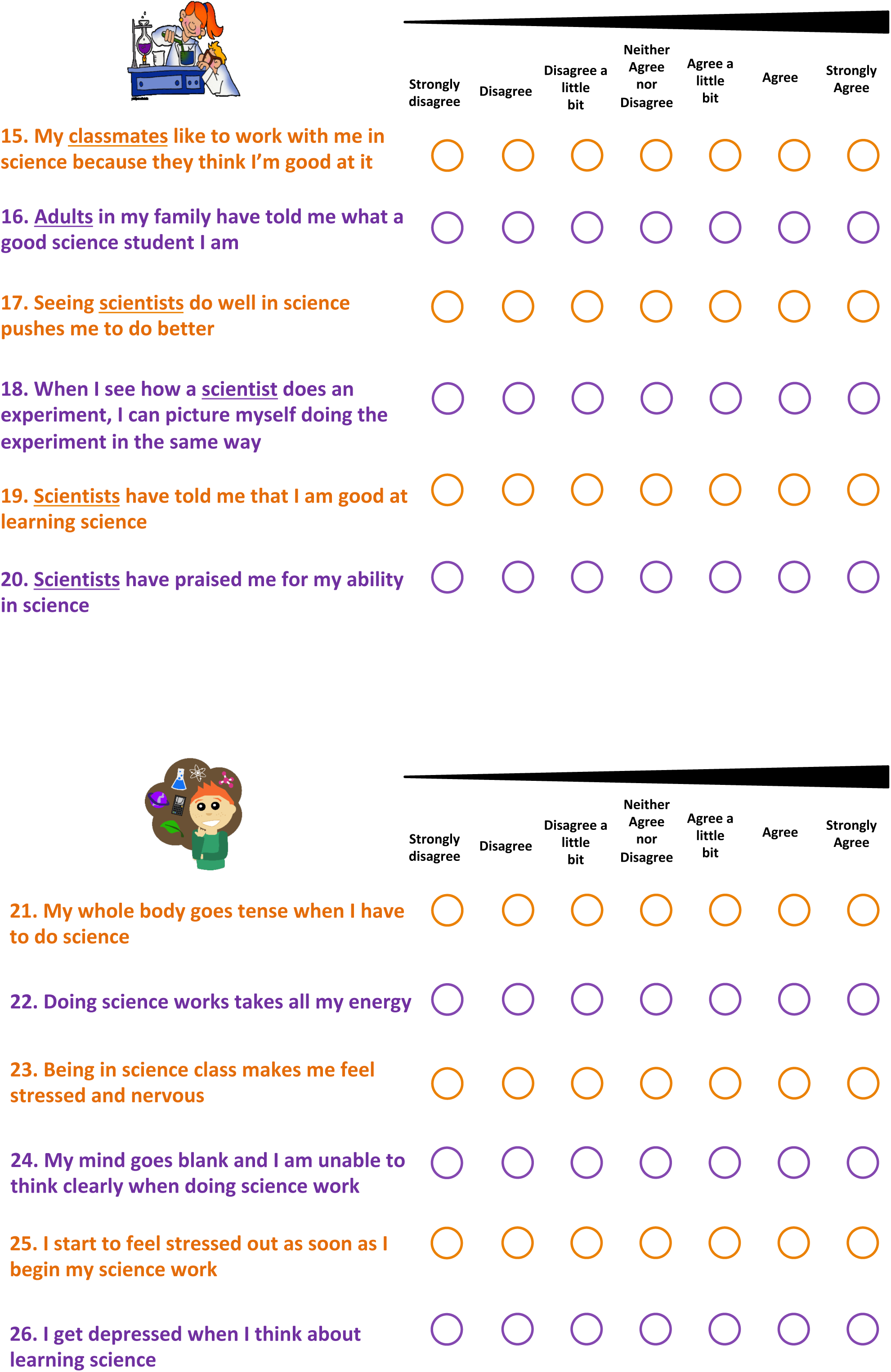
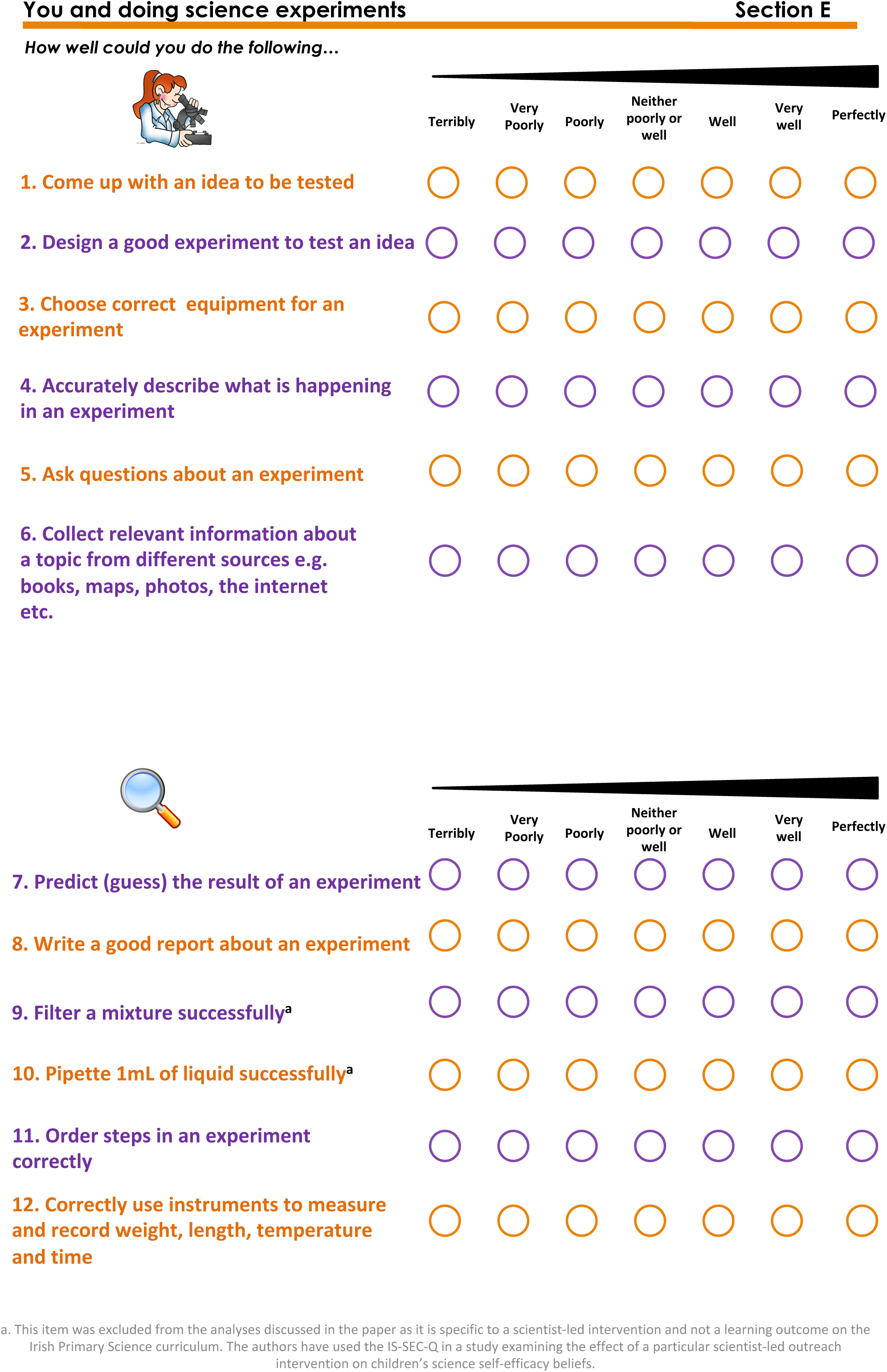

This questionnaire ends here! Thank you!

Please wait quietly until everyone else has finished.

Please remember to check over your answers.

Have you answered each question?

Check that you only have <u>ONE</u> answer per question

### Supporting Information

**S1 Table.** Coding scheme used for code interviews from Study 2.

| Research Question | Background/Justification for RQ | Interview Codes |
| --- | --- | --- |
| Do participants understand the report card section (section B)? | Possible misinterpretation/confusion as there was misinterpretations in the pilot | **Yes**<br>-Completed as expected<br>-Demonstrated understanding<br>**No**<br>-Not completed as expected<br>-Reasons for will emerge as sub-codes |
| Was there any topic in section C (Content knowledge) that participants didn't understand? | Checking to see whether topics from curriculum are comprehensible by children | **Yes**<br>What topic?<br>***Subcode***<br>*(specify)*<br>**No** |
| Was there any topic in section E (science skills) participants didn't understand? | Checking to see whether topics from curriculum are comprehensible by children | **Yes**<br>What topic?<br>***Subcode***<br>*(specify)*<br>**No** |
| How did children interpret D21-26 (emotional state)? | Items written for middle school children, quite negatively phrased. Some misinterpretations in pilot. | **As expected**<br>-Negative feelings<br>-Depressed<br>-Stressed/anxious<br>**Not as expected**<br>-Using physical energy<br>-Daydreaming |
| Was there any other source of confusion/comment children had? | Children were asked at the end of every questionnaire discussion if they'd anything else they wanted to add | **Yes**-other comment<br>**No**-no other comment |
| What kind of scientists were the children thinking about in D17-D20? | To see whether children were thinking of specific scientists | **Real**- specific real person, unspecific "real life" scientist<br>**Media**-Media, historical scientist, |

**S1 Table A.** Questionnaire comprehension

| **Research Question** | **Background/Justification for RQ** | **Interview Codes** |
| --- | --- | --- |
| Why did participants choose the middle option? | Issues in pilot: often picked as an “I don’t know option”, attempted to rectify by explaining in administration instructions that it was *not* middle option and instead reflects “average confidence” | **Average:** Middle confidence/average confidence<br>**Wasn’t sure:** or I don’t know<br>**Never happened** |
| Could participants differentiate between similar options in the scale? | It’s explained in the questionnaire administration instructions how the 7-point Likert-like scale works. Raised in literature that 7-point scale might be difficult for children. Points are labelled individually to combat this. | **Yes:** Gave rational<br>**Mixed:** Gave rational but doesn’t make sense/confusing<br>**No:** No rationale for choosing |
| How did participants rationale their choice of answer point? | To determine whether children are thinking about their item and can rationalise their answer | **No:** Could not give rationale<br>**Yes:** Gave rationale<br>**Yes Subcode** To emerge from data |

**S1 Table B.** Justifying response for choosing the middle option on the Likert-like response scale

| **Research Question** | **Background/Justification for RQ** | **Interview Codes** | **Questionnaire codes** |
| --- | --- | --- | --- |
| Does the answer for “Learn Science” in section A match with the participant’s answers in the interview to “Are you confident in science?” | Evidence for criterion-related validity | **Low confidence:** Not confident, not really<br>**Medium:** a bit confident, sorta confident<br>**High:** very confident, pretty confident | **Low:** 1-3<br>**Medium:** 4<br>**High:** 5-7 |

